# Cortical network mechanisms of anodal and cathodal transcranial direct current stimulation in awake primates

**DOI:** 10.1101/516260

**Authors:** Andrew R. Bogaard, Guillaume Lajoie, Hayley Boyd, Andrew Morse, Stavros Zanos, Eberhard E. Fetz

## Abstract

Transcranial direct current stimulation (tDCS) is a non-invasive neuromodulation technique that is widely used to stimulate the sensorimotor cortex, and yet the mechanism by which it influences the natural activity of cortical networks is still under debate. Here, we characterize the effects of anodal and cathodal tDCS on underlying neurons in active macaque sensorimotor cortex across a range of doses. We find changes in spike rates that are sensitive to both current intensity and polarity, behavioral state, and that are cell-type specific. At high currents, effects persist after the offset of stimulation, and the spatiotemporal activity associated with motor activity of the contralateral limb, measured by dynamics of neural ensembles, are altered. These data suggest that tDCS induces reproducible and noticeable changes in cortical neuron activity and support the theory that it affects brain activity through a combination of single neuron polarization and network interactions.

## Introduction

For nearly two decades, transcranial direct current stimulation (tDCS) has intrigued clinical and behavioral neuroscientists because it is simple to implement and there is evidence that it can produce clinical and behavioral gains. Most commonly, tDCS is delivered through two electrodes positioned on the scalp to deliver constant anodal or cathodal current over a target brain area for roughly twenty minutes. In comparison with other techniques for stimulating the nervous system, tDCS stands out because applied currents are not strong enough to directly induce firing in neurons.

Human tDCS is derived from “polarizing” stimulation that was delivered directly to the pia of motor cortex in anesthetized rodents^1–5^. The form of distributed stimulation was notable for its lasting influence over cortical excitability^2^ (i.e. excitation under the anode and inhibition under the cathode) that was later found to require NMDA-dependent plasticity^3,6^, along with changes at GABAergic terminals^7,8^ and BDNF release at glutamatergic synapses in motor cortex^9^.

Far less current reaches the brain during tDCS than during polarizating stimulation in animals due to technical and practical limitations^10–12^, but there are numerous clinical advantages. Most importantly, it is very safe and easy to apply to virtually anyone. And despite the differences in stimulation protocol, human experiments have shown lasting, polarity dependent, effects as well^13,14^. These seminal reports precipitated a huge number of tDCS studies over the past two decades, with experimental applications ranging from clinical rehabilitation (e.g. stroke^15^ and depression^16^) to basic electrophysiology (e.g. TMS-MEP^14^ and tDCS-EEG) and other physiological measures (e.g. magnetic resonance spectroscopy^7^), and behavior (e.g. reaction time and force production^17^).

In many cases, these new studies have raised even more questions about how tDCS works. It still is not clear how to best use it, nor has it become easier to predict its effect for untested conditions^15,18–20^. For example, while anodal and cathodal stimulation is generally considered to be excitatory and inhibitory respectively, this relationship appears to depend on duration and intensity^21,22^, brain state^23^, and other factors. Physiological and behavioral results are also variable across studies, but these differences are hard to interpret because these studies follow no standard methodology.

Considering the subthreshold nature of tDCS, a mechanism of action is hard to intuit and the responses of cortical neurons to clinical-type tDCS are unknown. *In-vitro* experiments suggest that, at magnitudes present in the human and monkey brain during tDCS^10,24–26^, electric fields can produce small (<1 mV) changes in the resting potential of the most polarized neuronal compartments^27,28^, which depends on morphology, orientation, and position of the soma^28,29^. Likewise, synaptic dynamics are also affected^30,31^. The cumulative effect of such factors is not straightforward, given the dense recurrent connectivity of cortex and its complex spatiotemporal dynamics.

To explain how weak subthreshold effects give rise to behavioral effects, a theory often called “amplification” proposes that network interactions between many simultaneously polarized neurons produces functional changes in brain spatiotemporal dynamics^32–34^. In fact, neuronal networks *in-vitro* are more sensitive to imposed fields than single neurons^35^. At the same time, some cast doubt on the idea that such weak intracortical stimulation produces meaningful effects, suggesting instead that tDCS activates peripheral afferents of the cranial nerves innervating the scalp^10,36^. The responses of individual neurons to tDCS during natural cortical processing are not well understood and remains a critical link towards settling this hypothesis.

Here, we characterize the effects tDCS on active sensorimotor cortical networks while monkeys alternated between coordinated forearm contractions and quiet sitting. We explore a range of clinically-feasible current intensities for anodal and cathodal tDCS while recording from overlapping populations of cells. Thus, we test the hypothesis that polarity impacts cell firing using both cross-over (by identifying neurons across experiments) and cohort analyses. We find that both polarities evoke a mixture of excitatory and inhibitory responses from the population, and a given neuron’s response to stimulation polarity is consistent across days. Furthermore, putative pyramidal cells are differentially affected by anodal versus cathodal stimulation, whereas putative non-pyramidal cells are less affected by tDCS overall, and have similar responses to both a- and c-tDCS.

We also measure the ensemble firing patterns reflected in population dynamics before, during, and after tDCS. While tDCS does not disrupt single neuron tuning to active wrist torques, we find that it alters how populations of cells coordinate their spiking activity during stereotyped movement. Overall, these results indicate that tDCS produces repeatable, modest changes in brain activity that results from polarization of the neurons.

## Results

### Changes in single-unit firing rates during tDCS

The most common type of tDCS targets sensorimotor cortex with one electrode, and places the other over the contralateral supraorbital area^14,37,38^ (**Figure 1a**). Human tDCS electrodes are roughly 5×7cm in size, and current densities are low (0.028 mA·cm^-2^ to ∼0.1 mA·cm^-2^)^39^. We delivered tDCS through saline-soaked cellulose sponge electrodes (3×3cm to accommodate for the smaller head size, **Fig. 1b**) and applied current densities that ranged from low in human studies (0.027 mA·cm^-2^, 0.25mA) to about four times that currently used (0.44 mA·cm^-2^; 4mA). These current densities are tolerated by patients (i.e. at the electrode/skin interface), but the intracranial manifestation of scalp stimulation varies between monkeys and humans, and even between human subjects. Thus, the precise mapping between stimulation amplitude and effect size in our study cannot be extrapolated to humans for numerous reasons (*Discussion* and *Supplemental Text*).

**Figure 1.**
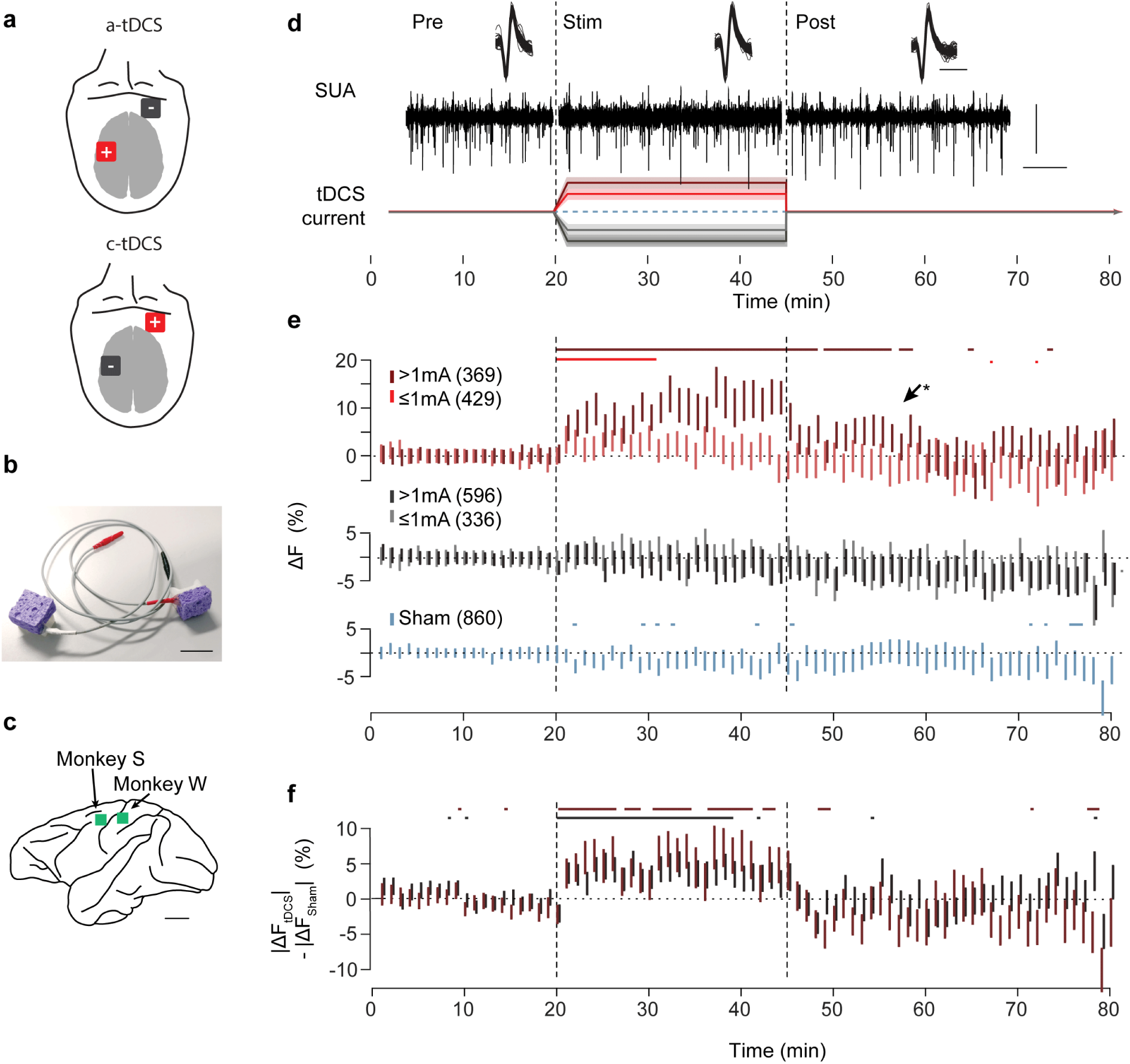
Simultaneous neural recording and clinical-type tDCS. **(a)** tDCS electrode montage for anodal or cathodal unilateral stimulation of sensorimotor cortex. **(b)** Cellulose sponge electrodes were designed to match those used in human trials. Scale bar: 3cm. **(c)** A microelectrode array was implanted on the gyral crown of left MI (monkey S) and left SI (Monkey W) in the areas corresponding to the right forearm. Scale bar: 1cm. **(d)** Time-course of experiment with example neural data. Top: sample SUA waveforms (scale bar: 0.5 ms) recorded during Pre, Stim (3mA a-tDCS), and Post epochs. Middle: continuous LFP filtered to spike band (scale bars: 100µV and 100ms). Bottom: experiment time-line. **(e)**. Time-course of *ΔF* (relative to average firing in Pre) during tDCS and Sham. The Stim/Sham epoch is indicated by two arrows and light gray background. Dark red: high dose (>1mA) a-tDCS, N=369; light red: low dose (<=1mA) a-tDCS, N=429; black bars: high dose c-tDCS, N:336; light gray bars: low dose c-tDCS, N:596; blue bars: Sham, N:860. **(f)**. Absolute changes in *ΔF* during high dose a- and c-tDCS is greater than during Sham stimulation. Vertical bars as in **e**, but calculated relative to Sham (|*ΔF*_tDCS_|*-*|*ΔF*_Sham_|). Vertical bars: median ± 95% CI, horizontal bars: p<0.01, Wilcoxon sign-rank test.

Throughout the experiments, monkeys performed a visuomotor target-tracking task by controlling the position of a cursor via isometric wrist torques registered by a 2-axis manipulandum (**Fig. 4a**). The task was intentionally simple and over-trained in order to maintain tight behavioral control and to isolate simple physiological changes in neuronal activity, and avoid confounding effects introduced by high-level cognitive processes (e.g. attention, learning). tDCS had little to no effect on the monkey’s ability to perform the task (**Supplementary Fig. 2**).

**Figure 2.**
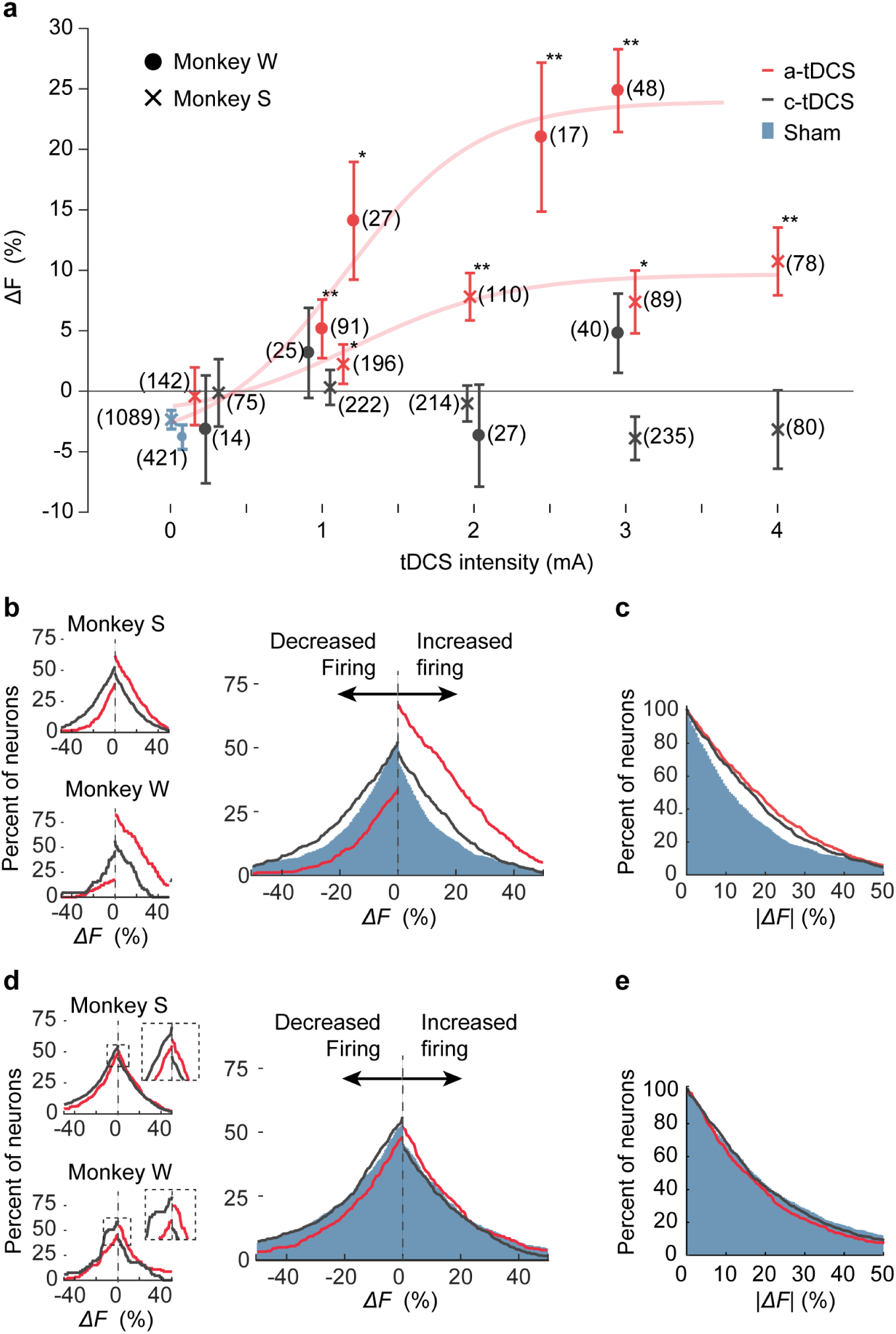
Firing rate changes depend on dose. **(a)** Change in firing rate (*ΔF*) between Pre/Stim for all recorded neurons. Average change in firing rate increased with dose during a-tDCS (red data points); the data is fitted with a sigmoidal dose-response curve *Supplementary Eq. 4* (data shows median ± 95% CI; a-tDCS: Monkey S: *x*_*50*_*=*1.20mA, *b*=0.85 percent·mA^-1^; Monkey W: *x*_*50*_*=*1.14mA, *b*=1.0 percent·mA^-1^). Effects of c-tDCS (black data points; Sham: blue data points) were masked by mixed changes in the population (two-sided Wilcoxon rank sum test tDCS vs. Sham, *p<0.01, **p<0.001 **(b,c)** Percent of cells with altered firing during tDCS or Sham. **(b)** During a-tDCS, the percent of neurons with increased firing was greater than sham (blue area), but the percent of neurons with decreased firing dropped. During c-tDCS, the percent of neurons with altered firing (both increased or decreased rates) was higher than sham. **(c)** Percent of neurons with absolute change in firing greater than or equal to |*ΔF*| was similar during a- and c-tDCS. **(d,e)** A similar bias in firing rate changes is still evident during the 30 minutes after tDCS was turned off **(d)** Percent of cells with altered firing after tDCS or Sham. Note that a-tDCS has more increases, while c-tDCS and Sham have more decreases. **(e)** Absolute size of firing rate changes is not greater than Sham during 30 minutes after tDCS is turned off.

**Figure 3.**
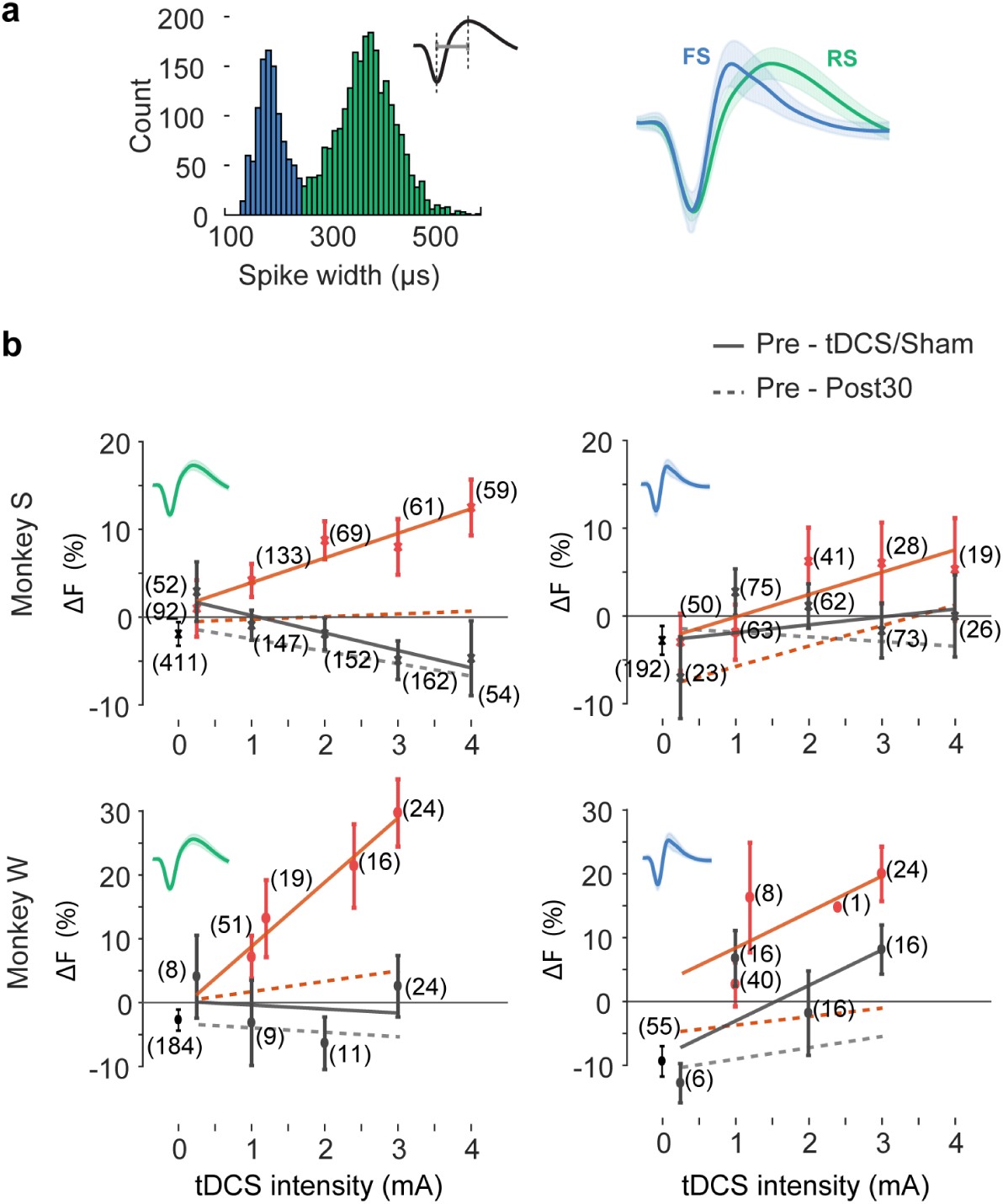
Putative pyramidal and non-pyramidal cells are differently affected by tDCS. **(a)** Histogram of spike width (peak-trough) for all neurons is bimodal and delineated two clusters (RS: ≥250µs, N=1812 and FS: <250µs, N=859). Right: average waveform of FS (blue) and RS (green) neurons (peak normalized for clarity). **(c)** Firing rates of RS cells are positivity correlated with a-tDCS dose and negatively correlated with c-tDCS dose (data points show median ± 95% CI. linear interpolation, *y=bx+c*. a-tDCS: Monkey S: b=2.8, R^2^=0.91; Monkey W: b=10, R^2^=0.96; c-tDCS: Monkey S: b=- 2.0, R^2^=0.87; Monkey W: b=-0.6, R^2^=0.02). Firing rates of FS cells tend to increase with a- and c-tDCS, and modulation is weaker than for RS cells (a-tDCS: Monkey S: b=2.5, R^2^=0.71; Monkey W: b=0.9, R^2^=0.13; c-tDCS: Monkey S: b=5.6, R^2^=0.52; Monkey W: b=5.5, R^2^=0.48). Dashed lines show linear interpolation of firing rate responses from Pre-Post30.

**Figure 4.**
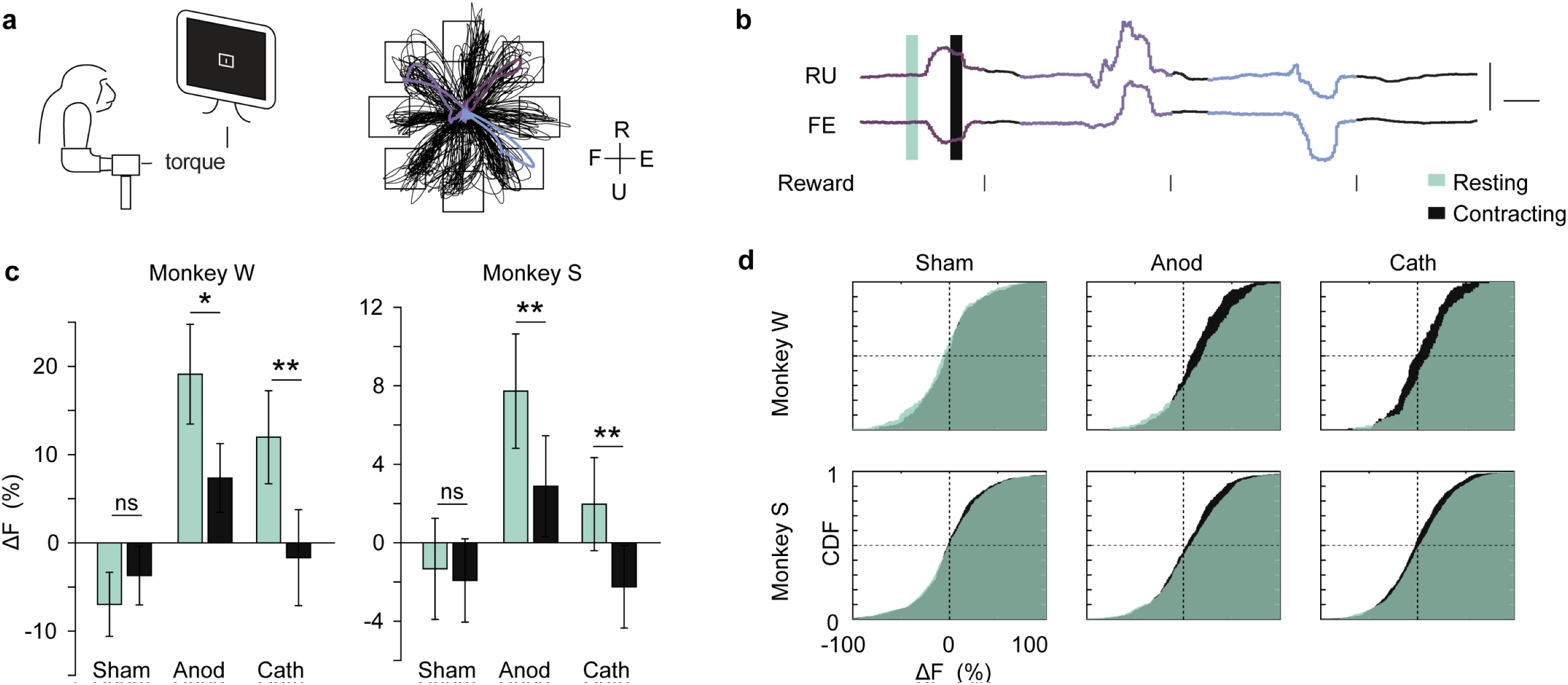
Firing rate responses vary between resting and contracting phases of task. **(a)** Task and example trials showing isometric torque in two dimensions. Torque trajectories (black traces) over 10 minutes of continuous task performance with target locations indicated by black squares. FE:flexion/extension RU:radial/ulnar **(b)** Three sample trials indicated by color in (a) for RU and FE torques. Each trial had Resting (before outer target cue) and Contracting (during isometric contraction) phases indicated by the tan and green boxes. Scale bars: 1N·m and 1 second. **(c,d)** Change in firing rate is different for Resting and Contracting phases of the task. **(c)** Median change in ΔF_pre,tDCS_ is the same for Resting and Contracting phases during Sham stimulation, but is significantly different for both a- and c-tDCS (all current doses). During a-tDCS, the greatest increases in firing rate are obvious during the Resting period. Error bars show ± 95% CI. **(d)** Cumulative distribution functions of the data underlying **(c)**.

Single unit activity (SUA) was recorded from the gyral crown by 96-channel microelectrode arrays (**Fig. 1c**) before, during, and after tDCS (**Fig. 1d**, Pre, tDCS, and Post epochs, respectively). We followed strict criteria for cell inclusion (*Methods: Single unit*), and over the course of 109 experiments (45 Sham, 29 a-tDCS, 35 c-tDCS) we obtained 2671 isolated unit recordings. The waveform and firing statistics of some of these units suggested that we recorded from the same neuron across days, so we used a cell identification algorithm to identify them (*Methods: Longitudinal cell analysis*) for cross-over analyses. Units isolated before the onset of tDCS were reliably recorded during stimulation (**Fig. 1d**). We tracked the proportional firing rate change of each neuron relative to its mean firing during the epoch before tDCS, 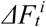 as 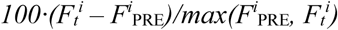 where *F*_*t*_ is the firing rate for neuron *i* at time *t* and *F*^*i*^_PRE_ is the mean firing rate during the “Pre” epoch. **Figure 1e** shows *ΔF* for all neurons during successive minutes throughout tDCS. Both high-dose (>1mA) and low-dose (≤1 mA) a-tDCS increased firing rates within the first minutes of stimulation. At high doses, increased firing persisted for about 15 minutes after tDCS was turned off (“*”, Wilcoxon rank-sum test p<0.01). Neither low-nor high-dose c-tDCS changed the population firing rates up or down, however, there was a similar *absolute* change in firing for both a- and c-tDCS as compared with Sham (**Fig. 1f**). This indicates that c-tDCS induced equal amounts of increased and decreased firing in the population.

The increase in population firing rates grew proportionally with applied current amplitude. As evident in **Figure 2a**, firing rates increased with increasing a-tDCS intensity, and the maximal sensitivity to tDCS intensity was similar for both monkeys (dose-response midpoint; S: 1.2mA; W: 1.14mA). On the other hand, c-tDCS produced no statistically significant increase or decrease in the population firing for any stimulation intensity. **Supplemental Fig. 3** shows these same results by experiment

### Percent of cells modulated by tDCS

Group statistics depicted in **Figure 2a** may mask mixed effects in the population, so we examined whether tDCS impacted the likelihood that any individual cell’s firing was increased or decreased. During a-tDCS, a higher percentage of neurons exhibited increased firing (*ΔF* >0), and a lower percentage showed decreased firing (*ΔF* <0) than during Sham or c-tDCS (**Fig. 2b**). Interestingly, a higher percentage of neurons had increased or decreased firing during high dose c-tDCS as compared with Sham, indicating that c-tDCS indeed affected firing, but the effects were mixed. To highlight this, **Figure 2c** shows the percent of cells with an absolute change in firing greater than or equal to a given |*ΔF*|. More cells had larger absolute firing rate changes during both a- and c-tDCS, and the percent of cells affected by c-tDCS is comparable to that affected by a-tDCS. Differences from Sham were more pronounced for larger absolute changes in firing – for instance, neurons were about 1.5x more likely to undergo a >20% change during a- and c-tDCS (corresponding to about 10-15% of neurons).

The bias in firing rate changes observed during tDCS (more increased firing than decreased firing during a-tDCS, and the opposite for c-tDCS) was still evident during the 30 minutes after it was turned off (**Fig. 2d**); however, the magnitude of these changes were no longer greater than those observed during Sham stimulation (**Fig. 2e**).

### Direction of modulation depends on cell type

Modeling and experiments *in-vitro* suggest that pyramidal cells are more susceptible to polarization than are symmetrical interneurons^27,28^, so we tested whether this was evident *in-vivo*. We used the width of extracellularly recorded action potentials to segregate putative pyramidal neurons from non-pyramidal neurons, a common analysis that was recently validated by cell-type specific optogenetic stimulation^40^ for these broad classes of neurons (but has limitations^41^). **Figure 3a** shows that the distribution of spike waveform width is bimodal, and the average spike shape of each cluster (pyramidal “RS” cells: ≥250µs, N=1812) and blue (non-pyramidal “FS” cells: <250µs, N=859). Consistent with other studies, FS firing rates were higher than that of RS firing rates, and exhibited task-related dynamics that were similar between monkeys (**Supplemental Fig. 4**)

Effects were evident in both cell types, but firing rate responses of RS neurons were larger and were sensitive to polarity. On the other hand, responses of FS cells were not sensitive to polarity. **Figure 3b** shows the spiking responses to a- and c-tDCS at different doses for RS and FS neurons separately. Note that firing rates of RS cells tend to get faster with increasing anodal currents, and slower with increasing cathodal currents. FS neurons are less correlated with tDCS intensity, but tend to increase firing in both conditions. Thus, population effects of c-tDCS are obscured by hetereogeneous responses related at least in part to cell type.

### Response during resting and contracting behavioral states

To determine whether the neural response to tDCS was different for the resting state versus active task performance, we extracted two periods from each trial (**Fig. 4b**, resting and contracting) and analyzed the change in firing rate from Pre-tDCS within these windows. **Figures 4c** shows that firing rate changes were more pronounced during the resting state than during active contractions. By contrast, firing decreased equally by a modest amount in both states during Sham stimulation. **Figure 4d** shows the complete distributions underlying the bar plots.

### The same neurons are consistently modulated by tDCS

We tested whether single unit changes were repeatable across sessions by identifying cells using a combination of cell identification methods^42–44^ (*Supplement: Longitudinal analysis*, **Supplemental Fig. 5-7**). The algorithm classified 1178 potentially unique neurons, 518 of which were recorded more than once (**Supplemental Fig. 6**). **Figure 5a** shows a typical example neuron recorded across four cathodal experiments, one sham experiment, and two a-tDCS experiments. The firing rate of this neuron decreased proportionally with c-tDCS intensity, but the pattern of firing in torque space (firing rate heat maps and directional polar plots) was preserved. For this neuron, firing rates remained constant during Sham and low-dose a-tDCS experiments (there were no high dose a-tDCS experiments with this neuron). We found that both a- and c-tDCS had significant repeat excitatory and inhibitory effects using a permutation test across all neurons recorded multiple times for a given condition (**Fig. 5c,d**, z score>5.3 or p<0.001, *Methods: Permutation test*). Neurons were about 2.5x more likely to undergo the same direction of firing rate changes during high dose a- and c-tDCS as compared with Sham (**Fig. 5d**, p<0.05), indicating that effects of tDCS are repeatable at the single cellular level. Furthermore, this analysis demonstrated that a-tDCS decreased firing in some neurons (**Fig. 5d**, *Always decrease*, z-score=5.3 or p<0.001), an effect that was not obvious from changes in firing rate changes alone. Neurons still showed a greater tendency to have consistently increased or decreased firing in the 30 minutes after tDCS was turned off (**Fig. 5c,d** “Pre-Post30”), although this effect was also more pronounced during Sham stimulation.

**Figure 5.**
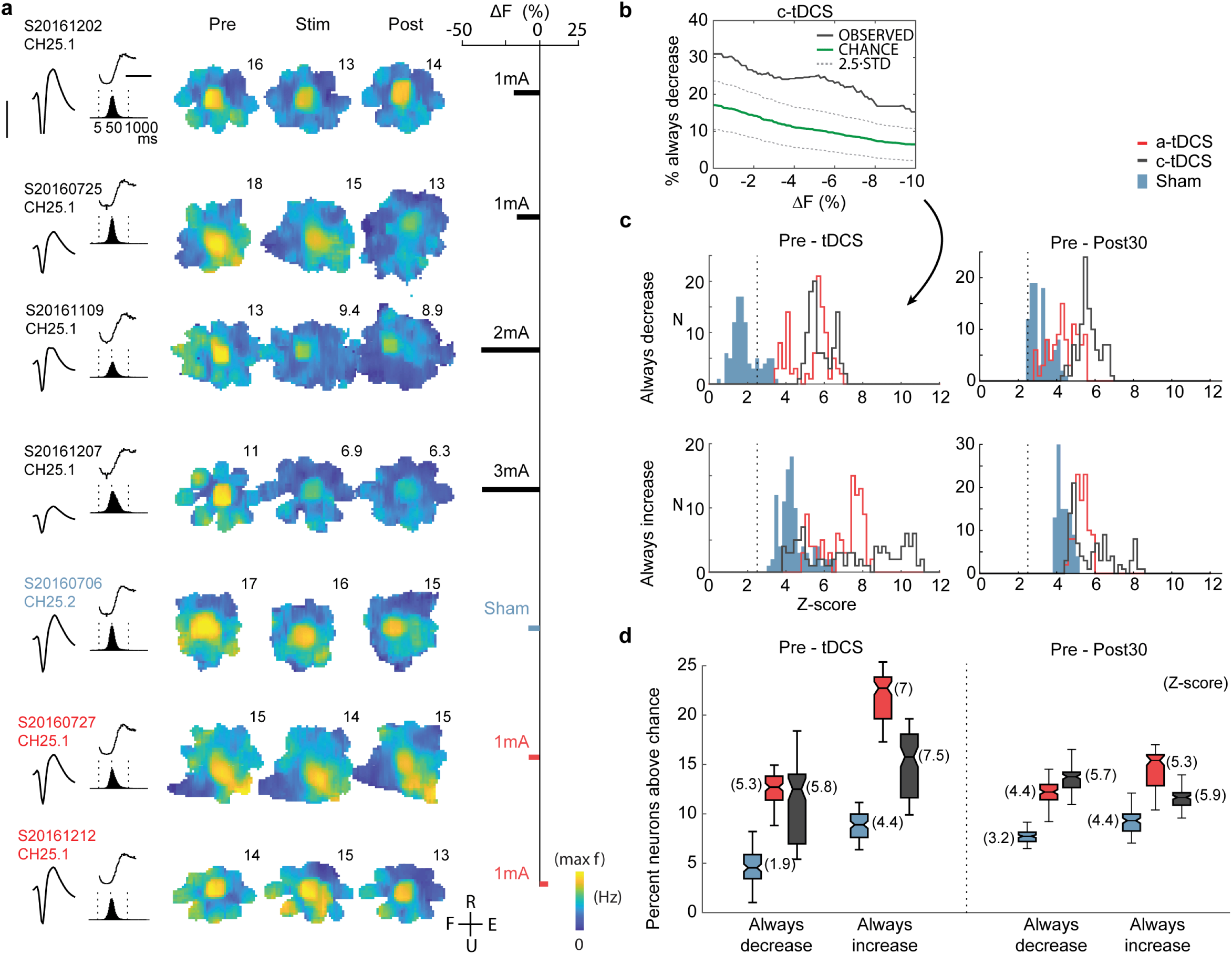
tDCS affects the same cells consistently across experiments. **(a)** A neuron recorded across 7 experiments (4 c-tDCS in black, 1 sham experiment in blue, and 2 a-tDCS experiments in red). From left to right, spiking statistics used to identify neurons across days: spike waveform (scale bar: 100µV), spike time autocorrelogram with lag [-20,100] ms (black trace, scale bar: 50ms), ISI distribution (black histogram), and firing rate maps in torque space for Pre epoch. Firing rate changes during Stim and Post epochs are evident in rate mapes, and and percent change in spiking from Pre-tDCS (bar). Cell firing rate drops with increasing c-tDCS intensity but is unaffected during sham or a-tDCS (no higher intensity a-tDCS experiments existed with this neuron). **(b)** We estimated the percent of neurons that consistently changed their firing by a given percentage within each condition by shuffling the observed *ΔF* 10,000 times. Here we show an example observed probability of consistent decreased firing during c-tDCS by at least *ΔF* from −10 to 0% (black trace). Chance probability ± 2.58·SD blue and dashed blue traces. **(c)** Distribution of z-scored probability that neurons are reliably excited or inhibited by tDCS. Dashed line indicates p=0.01 relative to data shuffled within each condition. **(d)** Percent of cells that always increased or decreased firing for a-tDCS, c-tDCS, or Sham experiments. Z-score relative to shuffled data indicated in parenthesis. Both a-tDCS and c-tDCS significantly increased the percent of cells that reliably increased or decreased their firing, suggesting that both polarities produce reliable, mixed effects in single neurons across sessions (z-scores>5.3, or p<0.001). During the 30 minutes after tDCS was turned off, neurons continued to exhibit more consistent changes than Sham. Box plot center line: median, notch: 95% CI, box: upper and lower quartiles, whiskers: 1.5x interquartile range (IQR).

### Single neuron tuning

Changes in spiking threshold can alter neuronal tuning by either sharpening or broadening the rate function^45,46^. We investigated whether tDCS disrupted neuron tuning to torque direction and torque space maps, which were prominent in both monkeys. To analyze a neuron’s preference for a given torque direction, we calculated the mean resultant vector (MRV) from the directional tuning function. The magnitude of the MRV (*R*_*L*_, *Supplementary Eq.* 6) describes the degree to which a neuron is directionally tuned, and its direction (*φ*_*R*_, *Supplementary Eq.* 7) indicates the preferred direction. Both measures are independent of firing rate and measure the shape of tuning only; therefore, firing rate changes induced by tDCS would not alone produce changes in *R*_*L*_ or *φ*_*R*_. **Figure 6a** shows the peak-normalized torque direction tuning curves for all recorded neurons ordered by *R*_*L*_ and rotated so that *φ*_*R*_ align. About one third of neurons recorded in each monkey were directionally tuned (*R*_*L*_>.1; Monkey S: 33.6%, Monkey W: 35.8%). **Figure 6b** shows five example cells with directional tuning which undergo firing rate changes during tDCS. While the amplitude of the tuning curves are altered, the shape is not, so *R*_*L*_ is not affected. This is clear from the population statistics shown in **Figure 6c**, which shows that tDCS did not induce changes in the direction or strength of tuning for any dose. Similarly, **Figure 6d** shows that the variability in directional tuning during tDCS was no greater than during Sham.

**Figure 6.**
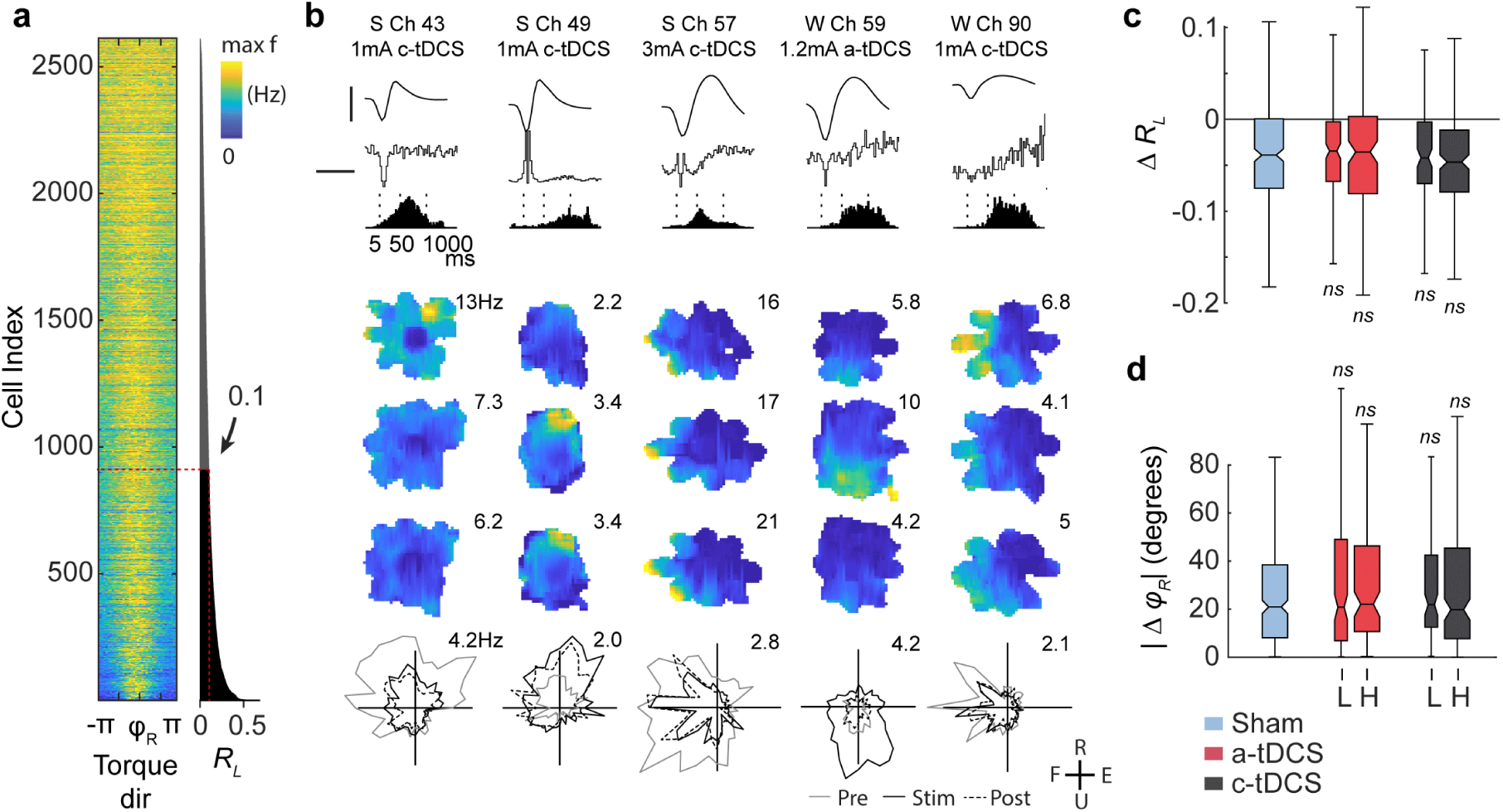
Amplitude but not shape of tuning is affected by tDCS **(a)** Directional tuning functions of all neurons (peak normalized and rotated by their preferred direction) are ordered by *R*_*L*_. **(b)** Five directionally tuned neurons (*R*_*L*_>0.1) during a- and c-tDCS. Spiking features from top to bottom: average spike waveform (scale bar: 100µV), spike time autocorrelogram with lag [-20,100]ms (black trace, scale bar: 50ms), ISI distribution (black histogram), firing rate maps in torque space for Pre, Stim, and Post epochs, and directional firing rate maps. While the amplitude of tuning is modulated by tDCS, directionally is preserved. **(c)** tDCS did not significantly change the strength of directional tuning as compared with Sham (L: low dose, ≤1mA, H: high dose, >1mA; p>0.05; Wilcoxon rank-sum test). There was a small but statistically significant decrease in *R*_*L*_ for all conditions (p<0.001; paired Wilcoxon sign-rank test). **(d)** Absolute change in preferred angle from Pre-Stim/Sham for directionally tuned neurons (*R*_*L*_>0.01). tDCS did not significantly change the direction of tuning relative to sham (|*Δφ*_*R*_| p>0.05), and there was no bias in the population-wide changes in preferred direction for any condition (*Δφ*_*R*_, not shown; p>0.05, Wilcoxon sign-rank test).

Beyond directional tuning, many neurons showed phasic task activation, such as increased firing during the resting versus contracting phases (see example neuron in **Fig. 5**). These correlations are reflected in the torque space firing rate map as hot spots, and we used these rate maps to assess cell tuning similarity (Pearson’s correlation coefficient, *ρ*) and specificity (bits·spike^-1^, *I*^47^) across epochs. **Figures 7a and c** explain how *ρ* and *I* vary with changes in the rate map. Cell tuning was relatively stable during the experiment (**Fig. 7b**, Sham), and high-dose tDCS produced a small but significant drop in *ρ* that reflected minor shifts in the rate map which exceeded those observed during Sham. Low-dose tDCS did not produce this change (**Fig. 7b**). Rate map shifts did not correspond with decreases in firing specificity (*I*) during tDCS experiments although they did during Sham (**Fig. 7d**). From these analyses we conclude that tDCS had very minor to no effect on single neuron tuning.

**Figure 7.**
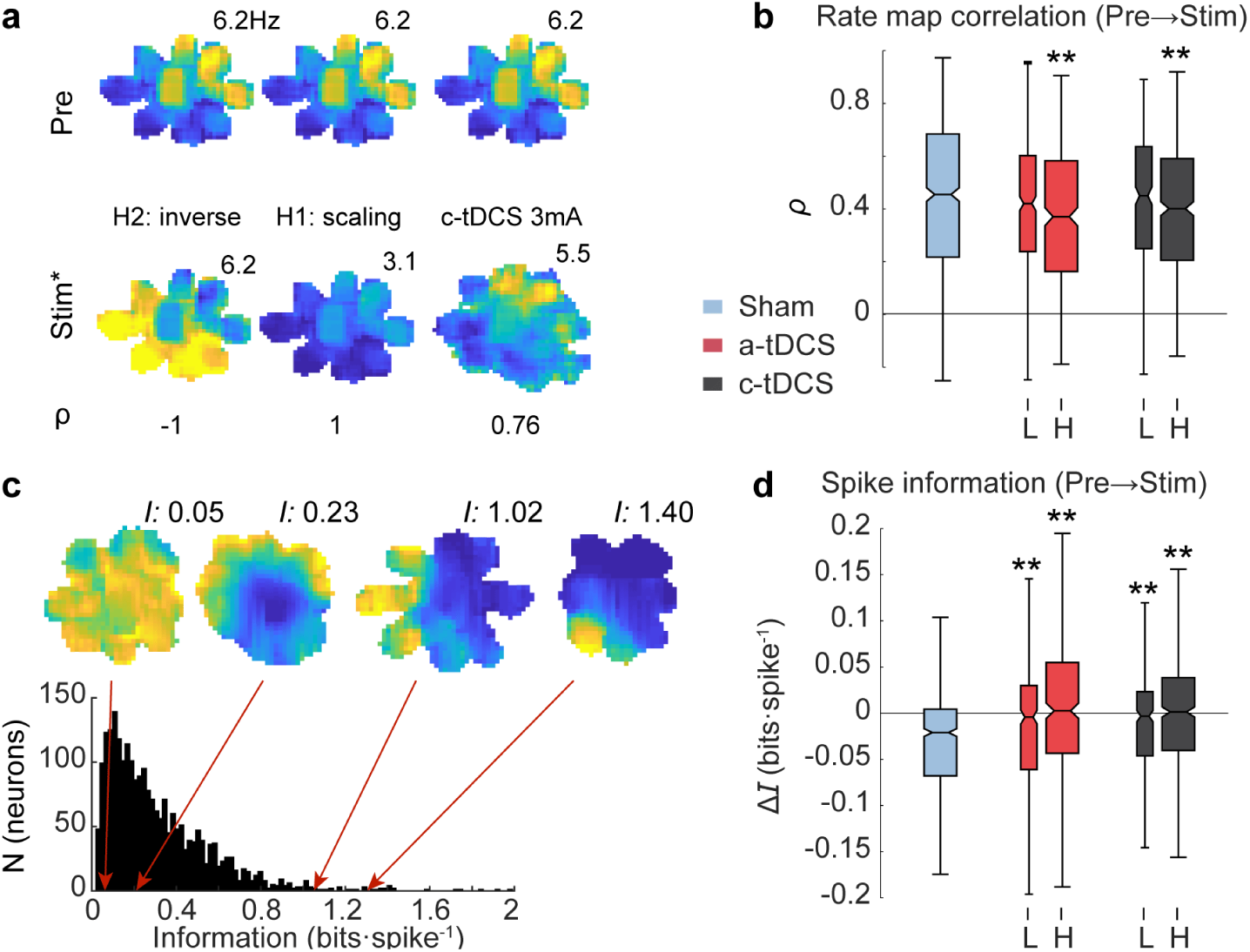
Small changes in torque space tuning during high dose tDCS. **(a)** Three rate map comparisons and their associated correlation coefficient (*ρ*) for illustrative purposes. ρ is minimal (−1) in the hypothetical case of inverted peaks/troughs, maximal (1) in the hypothetical case of single rate scaling, and high (0.76) for an example during during 3mA c-tDCS. **(b)** There were small shifts in tuning shape between Pre and Stim/Sham epochs, and this effect was more pronounced during high dose tDCS. (L: low dose, ≤1mA, H: high dose, >1mA) **(c)** Histogram of all torque vector spike information content (*I*) for all neurons. Four example rate maps with corresponding *I* values above demonstrate how “peaky” neurons carry more information than do neurons with dispersed spikes. **(d)** Shifts in tuning were not associated with drops in spike information during tDCS. (L: low dose, ≤1mA, H: high dose, >1mA; **p<0.001, Wilcoxon rank-sum test). Box plot center line: median, notch: 95% CI, box: upper and lower quartiles, whiskers: 1.5× IQR.

### Neural population dynamics

The expected influence of tDCS will be simultaneously present across synapses and neurons in a large patch of cortex, potentially affecting the correlations and relative timing of spiking between neurons. In this case, measurements of effects at the ensemble level could be more sensitive than those of individual neurons. To test whether (1) tDCS affected the size of functional neural ensembles, or (2) tDCS changed the spatiotemporal activation patterns of ensembles, we analyzed “neural dynamics”^48–51^ of simultaneously recorded neurons in a “neural space” whose dimensions corresponds to the firing of individual neurons^51–53^.

We analyzed smoothed spike trains (Gaussian filter σ=100ms) from 0 to 500ms after target appearance^52^ (this period contained neural activity most relevant to stereotyped contractions during the task), and applied a version of demixed principal component analysis (dPCA)^54,55^ to uncover the correlations in population firing. The averaged ensemble firing rates projected into the first two principle components resembled torque trajectories, and were well separated for the different targets (**Fig. 8a**). Since this plane reflected sensorimotor neural activity, we defined it as the ensemble “manifold”.

**Figure 8.**
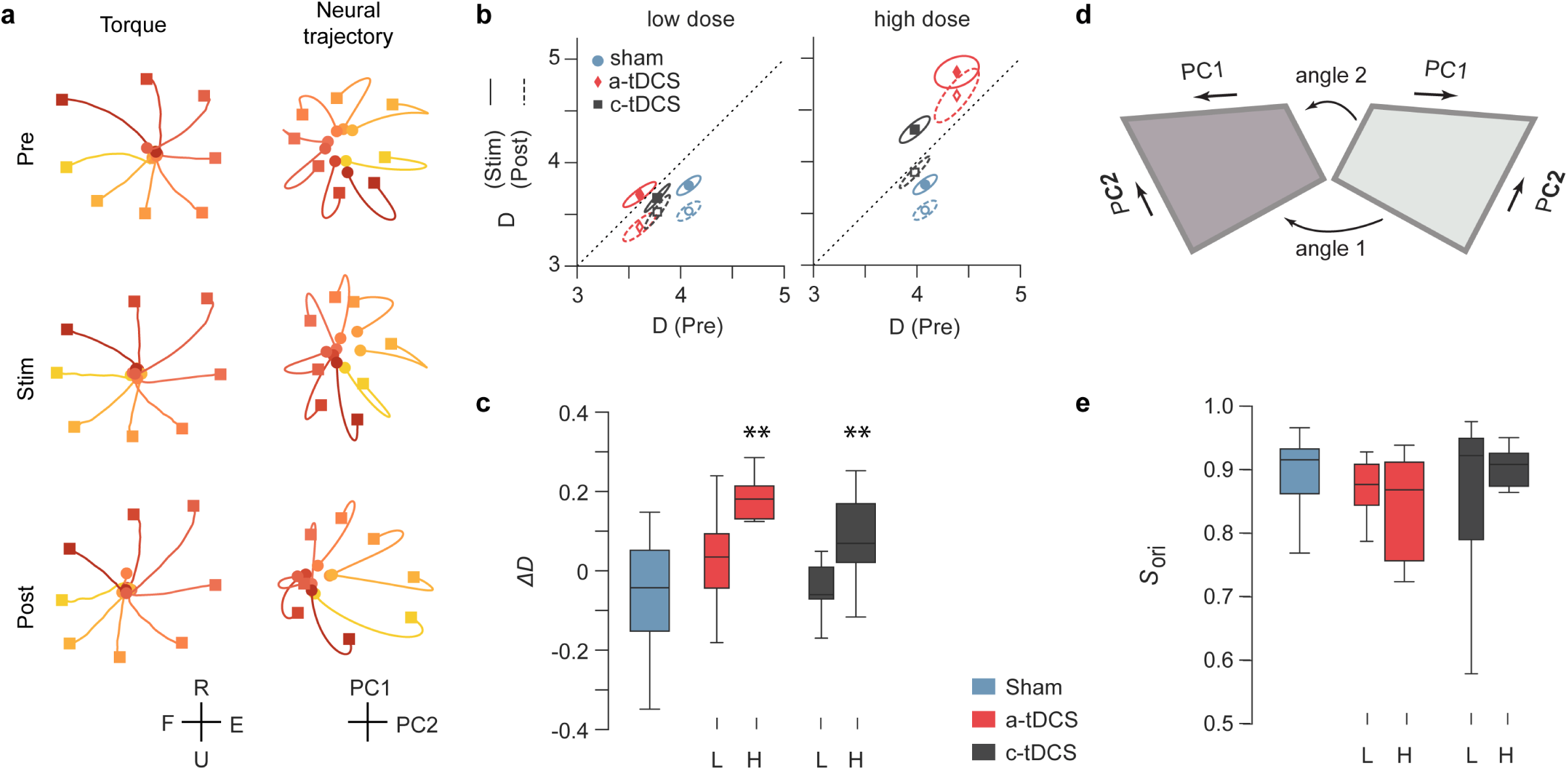
Population dynamics during the task and tDCS. **(a)** Target-grouped, averaged manipulandum torques (left) and population firing rates projected in first two PCs (right) for an example session, not normalized for illustration (3 mA c-tDCS, N_cells_=49). Colors indicate target identity, circle markers: t=0 seconds (target onset), square markers: t=0.5 seconds. **(b)** Mean dimensionality between Pre-Stim epochs (solid) and Pre-Post epochs (dashed). Points show mean of bivariate sample and ellipses show one standard error. Data points above the line (high dose tDCS during Stim and Post) indicate an increase in ensemble trajectory dimensionality, whereas points below the diagonal indicate a drop in dimensionality. Low dimensional ensemble trajectories utilize a smaller subspace of possible population patterns as compared with high dimensional trajectories. **(c)** Box plots showing statistics of Pre-normalized dimensionality change for neural trajectories during Pre and Stim. Box edges show first and third quartiles, internal bar shows mean, whiskers show extremal values. (L: low dose, ≤1mA, H: high dose, >1mA) **(d)** Illustration of principal angles between two linear PC manifolds. **(e)** Change in *orientation similarity (S*_*ori*_) of manifolds (PC1 & PC2 as depicted in **a)** from Pre to Stim. L: low dose, ≤1mA, H: high dose, >1mA. *S*_*ori*_ remains high during Sham epochs, indicating that ensemble patterns are stable over time. High dose a-tDCS evoked new dominant patterns of activity in the ensemble, indicated by a decrease in *S*_*ori*_. (*p<0.05, **p<.01, Sham vs. tDCS, independent t-test). Box plot center line: median, box: upper and lower quartiles, whiskers: 1.5× IQR.

We used two metrics to test whether ensemble firing changed during tDCS. First, we calculated the *dimensionality* (*D*) of the ensemble trajectories^56,57^. If all neuronal firing is highly correlated, points in the neural space approach a line (*D=1*), whereas if all neurons are firing independently, points are dispersed throughout the neural space (*D=N*_*cells*_). Population activity is typically restricted to a smaller subspace (1*<D<< N*_*cells*_), which is thought to reflect functional connectivity of the network^51^. In our experiments, *D* was similar across experiments (5.20±0.15). Interestingly, the dimensionality of ensemble trajectories decreased over the course of Sham experiments (**Fig. 8b,c**), whereas a- and c-tDCS both increased *D* (**Fig. 8b,c**, independent t-test Sham vs tDCS; a-tDCS p=0.007, c-tDCS p=0.008). Qualitatively, this result indicates that tDCS disturbed engrained ensemble dynamics and increased the working neural space during contractions. This effect was only significant for high doses, and was most pronounced during a-tDCS. Increases in ensemble dimensionality persisted after tDCS was turned off (**Supplementary Fig. 10**).

We also tested whether new dominant patterns of activity arose during tDCS by measuring the principal angles, *S*_*ori*_, between the intrinsic manifolds (PC1 & PC2) as schematized in **Figure 8d**. Manifold orientation changed very little during Sham epochs (*S*_*ori*_ = 0.89; **Fig. 8e**), indicating that ensemble activity patterns were stable over time. c-tDCS and low-dose a-tDCS did not produce shifts in the manifold, but there was a significant shift during high-dose a-tDCS (**Fig. 8e**; a-tDCS thick bars, p=0.045). Thus, high dose tDCS produced ensemble activity outside of the existing subspace, and a-tDCS was more effective at eliciting new dominant ensemble patterns than c-tDCS.

### Features in spike-triggered LFP are diminished during tDCS

Finally, we investigated the effects of tDCS on inhibitory synaptic currents correlated with neuron spiking observed in the LFP using the whitened spike-triggered LFP (wst-LFP)^58^. This method calculates the average spike-triggered LFP at multiple electrode sites and applies a spatial filter to distinguish the effects of a single neuron from non-specific components of the LFP which are common across the whole array (e.g., beta oscillations). Consistent with previous studies^58^, wst-LFP from our arrays support the notion that resultant features are synaptic currents evoked by spikes of the triggering neuron: they occurred after the spikes (**Fig. 9a,b**), and were restricted to electrode sites close to the neuron within known limits of axonal projection^59^ (**Fig. 9b,c**).

**Figure 9.**
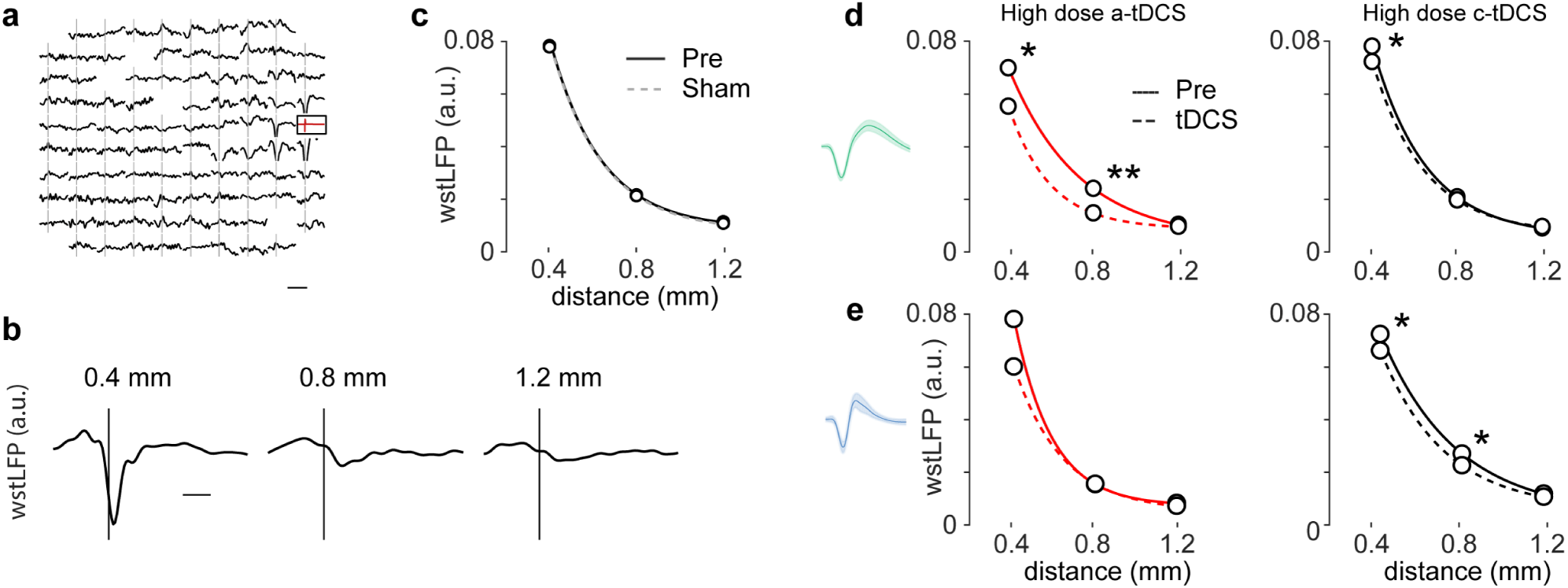
tDCS diminishes the amplitude of RS neuron spike-triggered LFP.**(a)** Whitened spike triggered LFP (wst-LFP) across the microelectrode array shows significant post-spike features at channels close to the neuron. Scale bar: 25ms. **(b)** Example wst-LFP at three distances from triggering neuron (Manhattan distance). Scale bar: 5ms. **(c)** Trough amplitude of wst-LFP (global minimum between 0 and 6ms lag) are stable from Pre to Sham epochs. The data is fitted with an exponential *A exp(-x/λ)+C*, where x is the distance and λ (0.24mm) is the space constant. **(d-e)**. Effects of high dose tDCS on RS and FS cells. (*N, A* and *λ* reported in *Supplemental table 1*). Effects did not persist after tDCS and are not shown (p>0.05). (**d)** Amplitude of wst-LFP at channels adjacent to RS cells is diminished during high dose a-tDCS (distance 0.4mm: p<0.01; 0.8mm: p<0.001; paired Wilcoxon sign-rank test). Effect is small, but significant for 0.4mm distance during c-tDCS (p<0.01). (**e)** Amplitude of wst-LFP at channels adjacent to FS cells is diminished during c-tDCS (distance 0.4mm and 0.8mm p<0.001) but not a- tDCS. **p<0.001, *p<0.01, paired Wilcoxon sign-rank test. In both cases, significant changes were not present after tDCS was turned off (p>0.05, not shown). See *Table 1* for fit parameters and corresponding N.

wst-LFP features were consistent throughout recordings without stimulation (**Fig. 9c**). During high dose a- and c-tDCS, however, there was a significant decrease in wst-LFP features for adjacent channels (0.4 and 0.8mm) when all cells were considered. Upon further inspection, we found that fields from RS cells were more affected by a-tDCS, while those from FS cells were more affected by c-tDCS. **Figures 9d,e** show how high dose tDCS affected RS (**d**, green spike) and FS (**e**, blue spike) neurons. Changes were restricted to neighboring electrode sites. Low-dose tDCS did not produce any significant changes (≤1mA, p>0.05). See *Supplemental Table 1* for fit parameters and corresponding N. We analyzed the wst-LFP features in the 30 minutes after tDCS was turned off, and found no persisting differences relative to Pre for any comparison (p>0.01, paired Wilcoxon sign-rank test).

**Table 1.** Fit parameters for Figure 7. The data is fitted with an exponential *A exp(-x/λ)+C*, where x is the distance and λ is the space constant. N_cells_ for this analysis are less than for others because some experiments were omitted due to poor LFP signal quality (N omitted, Sham: 8, a-tDCS: 10, c-tDCS: 15)

|  | High dose a-tDCS |  | High dose c-tDCS |  |
| --- | --- | --- | --- | --- |
| | $N$ | $\lambda_{pre}, \lambda_{post} (mm)$ | $N$ | $\lambda_{pre}, \lambda_{post} (mm)$ |
| All cells | 111 | 0.21, 0.21 | 324 | 0.28, 0.28 |
| FS | 38 | 0.18, 0.24 | 107 | 0.37, 0.31 |
| RS | 73 | 0.33, 0.19 | 217 | 0.25, 0.24 |

## Discussion

We recorded from populations of neurons in sensorimotor cortex during anodal and cathodal tDCS to determine if cells responded in a manner consistent with direct polarization. This data is important because it is difficult to infer the cumulative effects of distributed polarization on cortical neurons considering the dense connectivity of cortex, and its intrinsic dynamics during behavior. Neurons are susceptible to externally generated electric fields to varying degrees^27,28,34,35^, and synaptic transmission can be boosted or inhibited both pre- and post-synaptically^30,31^. This weak modulation will likely interact with natural brain dynamics in complex ways. For instance, although pyramidal cells may be more susceptible to direct polarization by tDCS, extrinsic excitation or inhibition of these cells could propagate through intermediate inhibitory cells to produce unpredictable network-level effects. Thus, characterizing the responses of neurons in normally active cortical circuits was one of the main goals of our study.

We detected firing rate changes in about 15% of neurons during tDCS as compared with Sham. Although the percent of cells modulated during a- and c-tDCS was similar, the proportion of neurons with increased firing was much higher during a-tDCS (9:1) than c-tDCS (3:2). The hetereogeneous cellular response to cathodal stimulation may help explain the fact that c-tDCS sometimes produces excitatory effects in human studies. Importantly, the same percentage of cells, 15%, was found when estimating the percent of neurons that reliably modulated their firing in the same direction for repeat a- or c-tDCS. Overall, despite day-to-day differences in the composition of neurons sampled, the average response to a given polarity and dose was very repeatable across days for individual neurons (**Supplemental Fig. 3**); not only were the average population statistics similar across days, single neurons were also more likely to be consistently modulated up or down by high dose tDCS.

It is understandable that firing rate changes were only detectable in a subset of neurons. Neurons differ in their response to polarizing currents, and even in those most polarizable, various cellular compartments may undergo a very modest (<1mV) change in membrane potential. While some network models have predicted that such changes could produce changes in spike timing^33,45^, it was not clear whether tDCS would produce changes in firing rates. In fact, an earlier study did not detect firing rate changes^60^, perhaps due to the smaller sampling of neurons, or because only one lower intensity current was applied for shorter time intervals. We also found that firing rate responses were greater during the resting state as compared with the contracting phase of the task. This could be due to the fact that an increase in spiking probability will be more pronounced during periods of relative quiescence.

Our recordings came from a depth of 1.5 mm at the gyral crown, directly under the stimulating electrode where the electric fields are strongest^10,25^ and are expected to be radial to the cortical surface. The responses of putative pyramidal cells changed with tDCS intensity and polarity – increasing with anodal stimulation and decreasing with cathodal stimulation. These findings are consistent with the “somatic” theory of tDCS, and are suggestive of direct polarization of the neurons following the pattern of hyperpolarization and depolarization expected by morphology and orientation in the field^27,28,34,35^. In contrast, the firing rates of putative inhibitory cells positively correlated with tDCS intensity regardless of polarity, which may be the result of dendritic depolarization, or secondary to network effects following the primary modulation of other cell types. Errors induced by this method of cell identification^41^ could lead to an underestimation of cell-type dependency on polarity, so this effect may be larger than we could detect.

Neuronal tuning is often compared to an “iceberg”^45,46^, in that a relative shift in spiking threshold may change the shape of the tuning curve: causing it to become more dispersed when the soma is depolarized, and sharper when it is hyperpolarized. However, we did not observe obvious changes in the broadness or direction of single unit tuning, indicating that tDCS did not simply increase excitability at the expense of selectivity. Effects were similar in both pre- and post-central gyrus, which have analogous structures and overlapping functions^61–63^. In both cases, tuning to torque was preserved, which presumably reflect a mixture of both efferent and afferent cortical signals and suggests that responses in other cortical areas with analogous cell organization could be similar.

We analyzed neural ensemble dynamics to measure changes in how neurons fire relative to one another. Ensemble dynamics are confined to a small part of the total neural space^51^ (the manifold) and the shape of this activity is probably determined by functional connectivity patterns in the network^51,64^. In our experiments, dimensionality of ensemble trajectories increased during tDCS, although it tended to *decrease* with time otherwise. This finding is consistent with theories underlying the warm-up effect, in which a network refines its activity during practice of a rehearsed task^65^. Conversely, increasing the dimensionality (and therefore dispersion) of firing during tDCS may explain tDCS-associated increases in learning rates: by increasing the size of the working neural subspace, new, successful patterns can be more quickly tried and reinforced^66^. Therefore, it may be that high dose tDCS releases constraints imposed by functional connectivity patterns (by dendritic depolarization, disinhibition, etc.) to augment learning from otherwise deeply embedded associations.

tDCS is predicted to affect synaptic transmission^30,31^, so we tested this by measuring features of the unitary LFP using the wst-LFP. Deflections in the wst-LFP are thought to reflect inhibitory postsynaptic potentials in apical dendrites of pyramidal cells^58^. tDCS decreased the amplitude of these currents, an effect most pronounced among RS cells during a-tDCS, which coincided with a global increase in firing. This is consistent with direct modulation of RS cells by tDCS with a consequent drop in feedback inhibition and decreased inhibitory drive while the apical dendrites are predicted to be hyperpolarized. Also, the larger decrease in wst-LFP associated with RS, but not FS, cells could be due to divergent effects resulting from documented connectivity patterns from RS neuron to many inhibitory interneurons^67^. Decreases in RS firing during c-tDCS did not correspond with an increase in wst-LFP, suggesting that c-tDCS directly inhibited RS cells. One puzzling result is the decrease in wst-LFP associated with FS cells during c-tDCS, a condition when the apical dendrites of pyramidal cells are predicted to be relatively depolarized. Future studies in vitro may elaborate on this interplay between simultaneously polarized synapses and neurons.

There was a notable difference in firing rate responses between the two monkeys used in our study, which may be due to inter-subject anatomical differences. In fact, Opitz et al., found a two-fold difference in electric field magnitudes between two monkeys, with more current reaching the brain in the smaller one^25^. Similarly, there was a sizeable difference in weight and head shape between the monkeys in our project as well: monkey S was significantly larger and had more epicranial muscle mass than monkey W (14kg versus 10kg). Firing rate changes in the larger monkey were smaller than those observed in the smaller one (**Figure 2a**), suggesting that these anatomical differences played a role.

Compared to any other animal model for studying motor behaviors and neuromodulation of motor cortex, non-human primates are the most biofidelic. At the same time, model artifacts exist: the brains of monkeys are smaller and with less cortical folding than humans; the shape of the head may impact the way current flows between the stimulating pads; differences in neuronal phenotypes could influence the response properties; and implanted devices for intracortical recording can also alter the way current flows through the skull. Of these, a particular focus in the field has been the magnitude of the intracranial electric field – and whether currents applied to the scalp are a reasonable metric for dose. Fortunately, many studies have now measured the electric fields induced by scalp stimulation in humans and monkeys, and suggest that the field magnitudes are very similar. In monkeys, intracranial recordings found that 1mA tDCS delivered through 3.14cm^2^ electrodes (0.32mA/cm^2^) resulted in median field of 0.2 and 0.4V·m^-1^ in two animals^25^, and another study found that 2mA scalp stimulation through 10.2cm^2^ electrodes (0.2mA/cm^2^) induced 0.12V/m fields^24^. Likewise, in human studies, 2mA delivered through 4cm^2^ electrodes (0.5mA/cm^2^) produced 0.8V/m fields^26^, and 1mA stimulation through 25cm^2^ (0.04mA/cm^2^) electrodes produced about 0.2V/m^25^. Current density is a reasonable surrogate for electric field across these studies, which used both monkeys and humans, different electrode shapes, sizes, and montages, and currents - and which found results that were similar within an order of magnitude. We explored a 16-fold range of current densities in our study, so it is very likely our stimulation protocol overlaps with field magnitudes induced by human tDCS.

Some have proposed that effects of transcranial stimulation are mediated by afferents of the cranial nerves innervating the scalp^10,36^, rather than by direct effects on brain cells. Recordings from both monkeys were collected from an area of cortex innervating and innervated primarily by the contralateral forelimb, so it is unlikely that observed changed were due to activation of the trigeminal nerve. Nevertheless, sensory disturbances could have caused behavioral changes that would be reflected in motor cortex. We do not believe that the changes in spiking we observed are due to such peripheral effects for several reasons. First, the monkeys did not appear to notice the stimulation, and continued to perform the task at rates/torques nearly identical to pre-stimulation epochs (**Supplemental Fig. 2**). Monkeys are very sensitive to such changes, and sensory or visual distractions would have likely resulted in marked behavioral changes. Second, tingling or burning sensations would be similar for anodal and cathodal stimulation, and in this hypothetical case, alterations brain activity would not be different for the two conditions. However, this was not the case: responses evoked by anodal and cathodal stimulation were significantly different. First, the proportion of neurons excited and inhibited varied, and second, the pattern of excitation or inhibition in single neurons matched the expected patterns given by their cell type.

The clinical utility of tDCS depends on additional physiological data that will refine the protocol and further test the underlying mechanism of action. Many believe that intracranial current from tDCS directly polarizes neurons in the brain, but this has been called into question^10,36^. To address this, we examined the patterns of modulation across populations of cortical neurons and found effects consistent with direct neuronal modulation during anodal and cathodal tDCS at high current intensities. Quantification of ensemble dynamics demonstrated that anodal tDCS was more effective at shifting network activation patterns, but both polarities expanded the active neural space and had effects that outlasted stimulation. Thus, at sufficient doses, it appears that tDCS acts on underlying cortex through a combination of cell polarization and second-order network mediated interactions. Future studies could investigate the potential for these changes to beneficially affect learning and plasticity in the behaving primate.

## Methods

### Subjects and behavioral task

All experiments were conducted with two male Macaca nemestrina monkeys (S and W) and conformed to the National Institutes of Health ‘Guide for the Care and Use of Laboratory Animals’. Procedures were approved by the University of Washington Institutional Animal Care and Use Committee.

Monkeys were trained in a visuomotor target tracking task conducted in a primate behavior booth equipped with a computer monitor (30 cm × 23 cm), tones for audio feedback, and a computer-controlled feeder dispensing fruit sauce as a reward. Monkeys voluntarily moved from the cage to a primate chair which was adjusted to allow each monkey to sit upright comfortably. The left arm was loosely restrained in a plastic tube, and the monkey used an isometric manipulandum with the right arm (contralateral to the recording array) to control the position of cursor on the screen using torque registered in two dimensions (horizontal axis: flexion/extension, FE; vertical: radial/ulnar, RU). The head was restrained with flexible plastic bars for the duration of experiments to promote attention in the task and for stability of neural recordings.

Each trial consisted of four phases: OFF, START, CONTRACT, and RELAX. At the beginning of a trial, the screen was blank for five seconds (OFF), followed by a cue (START) that indicated the monkey would soon have to produce a torque about the wrist to one of eight possible targets (CONTRACT). During the START condition, the monkeys had to keep the cursor in a center target (no contraction) for a variable period of time (0.5-2 seconds) so that we could measure response time. During the CONTRACT condition, targets pseudo-randomly selected from one of eight targets in FE and RU were presented. After holding the cursor in the CONTRACT target box for 1 second, the RELAX target would appear in the center of the screen, cueing the monkey to relax his wrist to receive a reward. The monkeys performed the task continuously for the entire duration of the experiment, and each experiment consisted of roughly 1000 trials.

### Surgery and implants

Both monkeys underwent a two-stage implantation schedule. For all surgical procedures, the skin around the surgical site was shaved and scrubbed with betadine. Sterile surgery was performed with the animal under 1–1.5% isoflurane anesthesia (95:5 O2:CO2). Cardiac waveform, heart rate, respiratory frequency, blood-pressure and end-tidal CO_2_ was monitored continuously. Post-operative care included protected recovery in a cage, administration of analgesics (ketoprofen 5 mg/kg), additional doses of antibiotics (cephalexin 25 mg kg-1 PO), and careful overnight observation by trained personnel. After recovery, monkeys showed no signs of discomfort related to any of the implanted devices.

During the first surgery, four 1.25×4mm perforated titanium craniofacial ties were affixed to the skull by 3 2×6mm titanium bone screws each. Two were implanted bilaterally over the occipital ridge, and two were placed temporally bilaterally. There was an interval of >6 weeks before the next surgery to allow the plates to osseointegrate with the skull.

During the second surgery, a 96-channel microelectrode array (length = 1.5mm, Blackrock Microsystems, Salt Lake City, UT) was implanted (left MI, Monkey S; left SI, monkey W) using the pneumatic insertion technique^68^. To implant the array, a bone flap (2×2.5cm) was removed from an area of the skull located stereotactically. Electrode arrays were placed in hand/wrist sites in pre and postcentral gyrus, located in precentral gyrus from the line extending from the genu of the arcuate sulcus posteriorly to the central sulcus (Monkey S), or in the adjacent location in postcentral gyrus (Monkey W). After placement of the array, the dura was reapproximated over the array and the bone flap was replaced in the craniotomy with two 1mm screws and a single strip of thin titanium mesh plate together with hydroxyapatite to promote skull regrowth around the margin of the bone flap. The skull was inspected post-mortem (Monkey W) at the end of experiments, and there were no remaining defects or holes in the skull except for the hole permitting passage of the wire bundle. One connector pedestal was fixed to the skull caudal to ear-bar-zero. Finally, a lightweight aluminum halo system used for head fixation and affixing the tDCS pads was mounted with four pins seated in each of the four bone straps implanted during the first surgery. These pins were low on the head and separated by >2 cm from the tDCS pads. Daily recording sessions began after the monkey had completely recovered from surgery.

### Transcranial direct current stimulation

tDCS was designed to match human trials as closely as possible. Scalp pads were made from cellulose sponges soaked in 0.9% NaCl solution with inlaid copper wire, and were cut to a smaller size to accommodate the smaller anatomy of the monkey brain and skull (**Fig. 1b**; 3×3cm versus ∼5×7cm. Sponges were checked so as to be moist throughout but not wet or dripping onto the scalp to avoid any undesired current spread. Similar to the classic sensorimotor cortex/contralateral supraorbital montage used in human trials^38,69^, we placed one pad over the implanted array in sensorimotor cortex (anode: a-tDCS, cathode:c-tDCS), and a second pad over the right supraorbital ridge (main text **Fig. 1a**). Proper location of pads was easily determined relative to landmarks (halo, pins, and connector pedestal) each day. As necessary, hair was removed with Nair™ (Church & Dwight Co.), and the skin was rinsed and dried. At the onset of tDCS, current was delivered by a manual stimulator as previously used in clinical trials (Phoresor II autostimulator, IOMED, Salt Lake City, UT). This stimulator sounds an alarm and aborts stimulation if the compliance voltage (60V) is reached during stimulation. At 4mA, this corresponds to an electrode/tissue impedance of 15kΩ, which is well above our normal impedances (∼5kΩ) and those reported during human tDCS. We tested the stimulator’s ability to deliver 25 minutes of constant stimulation near its compliance voltage (**Supplemental Figure 1**). tDCS was slowly ramped up over 1 minute to a predetermined current to decrease the chance of the animal perceiving the stimulation. If noise was observed in the neural recording as tDCS current was ramped, stimulation was decreased to the nearest dose that did not affect recording (3 experiments). Similar to human trials^69^, the duration of tDCS was 25 minutes, and we explored doses ranging from the lowest delivered to human subjects (0.027 mA·cm^-2^) to roughly four times the highest dose currently delivered to humans (0.44 mA·cm^-2^).

#### Experiment time course and behavioral task

During each experiment, the monkeys were transported from their home cage to the recording booth in a primate chair. Once in the booth, a feeder tube was placed in front of the monkey’s mouth to deliver fruit sauce reward. Finally, we prepared the scalp and placed the tDCS pads (see **tDCS**, above), and connected the recording system to the microelectrode array.

At the start of the experiment, we initialized the behavioral task (custom MATLAB software), which ran without interruption until the experiment ended. Epoch 1 (Pre) lasted 20 minutes, and established baseline activity. Epoch 2 (Stim/Sham) lasted for 25 minutes, and had three possible conditions: a-tDCS, c-tDCS, or Sham. Epoch 3 (Post), lasted over 30 minutes, or until the monkey showed signs of fatigue or disinterest in the task. The monkeys often performed the task for up to an hour following tDCS.

### In vivo neural recordings, task and behavioral data

We used a 256 channel digital data acquisition system (Tucker-Davis Technologies) to record neural, behavioral, and task data in the primate booth. Data from the manipulandum (isometric torque) and behavioral task (target number, cursor position, task condition) was sampled at 3kHz. Voltage signals from the microelectrodes were amplified, digitized, and streamed to the TDT base station with a sampling rate of 25kHz. The data streams from each electrode were processed and saved as local field potential (LFP) and single unit activity (SUA). To extract the LFP, signals were band-pass filtered (0.25 – 500 Hz), downsampled (3 kHz), and saved to hard disk. To extract SUA, signals were band-passed filtered (0.5-4 kHz), and a fixed threshold unique to each channel was used to detect action potentials by negative threshold crossing. Thresholds were initially estimated automatically as minus three times the standard deviation estimated from 10 seconds of the SUA filtered signal, and the first five minutes of recording was used to ensure that thresholds were appropriately set outside of the baseline noise and to detect spikes. When a spike was detected, a snippet of SUA signal was recorded to the disk corresponding to 8 samples before the threshold crossing and 22 samples after the threshold crossing (1.2ms window)

### Single unit identification and inclusion

Snippets of detected spikes in the SUA signal were sorted offline with Offline Sorter v4 (Plexon Co, Dallas, TX) using the template matching method. For each neuron, the signal to noise ratio (spike SNR)^70^ was calculated by equation 1:

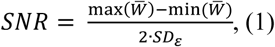

where *W* the collection of all spike waveforms, 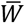 is the average waveform, and ε is the matrix of noise values calculated as deviations from the mean:

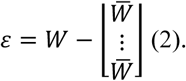

Only neurons with >1000 spikes, an average peak height of >50uV, and spike SNR >3 were included for analysis.

### Statistical analysis

All statistical comparisons were calculated using MATLAB R2017a or Python Pandas module. When applicable, we used paired statistical tests and did not assume normality. In particular, Wilcoxon’s paired, two-sided sign rank test (signrank, MATLAB) was used when comparing cells across conditions, and the Wilcoxon’s two sided rank sum test (ranksum, MATLAB) was used when comparing the behavioral and neural activity across sessions. Unless otherwise noted, data and error bars depict the median and the 95% confidence interval of the median, respectively.

#### Changes in firing rate

Data depicted in main text **Figure 2** was generated by compiling data across multiple experiments for each current step. The data from each individual experiment can be seen in **Supplemental Figure 3**. For these analyses, we calculated *ΔF*, which describes the percent change in firing between Pre and Stim, and was calculated by equation 3:

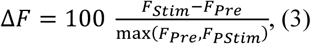

where *F*_Pre_ and *F*_Stim_ are the median firing rates for the Pre and Stim/Sham epochs. Equation 1 is bound between −100 and 100.

We fit a sigmoid function to the data presented in **Figure 2a**, defined by equation 4:

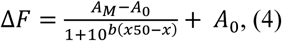

where A_0_ is a fixed parameter defined by *ΔF* during Sham epochs for each monkey, and *A*_*M*_, *x50*, and *b* are free parameters describing the maximal value, inflection point and inflection point slope, respectively.

### Longitudinal cell identification

The microelectrode arrays used in these experiments are chronic, immovable, and can record stable SUA for years^71^. Therefore, it is likely that some neurons were recorded across many days. We leveraged this to investigate whether a-tDCS or c-tDCS reliably affected individual neurons across experiments. We performed an analysis inspired by previously reported work^42–44^ to identify which spikes might originate from the same neuron across days using various spiking statistics (**Supplemental Figure 6)**. First, we calculated five metrics for each neuron: the mean waveform, spike rate during Pre, refractory period-timescale autocorrelation (bins logarithmically spaced from decades 10^−3.6889^ to 10^-1^ms), regular-timescale autocorrelation (bins linearly spaced from 0 to 100ms), and ISI distribution (bins logarithmically spaced from decades 10^-3^ to 10^1.5^ms). Logarithmic spacing was used for short-timescale autocorrelation and ISI histogram so that the shape of short time interval features (where many important dynamics are reflected) were weighted similarly to slower ones.

The distance between each of the five metrics was calculated pairwise across all neurons. The L^2^ norm was used for ISI distribution, autocorrelation, and firing rate, whereas the distance between spike waveforms was defined as 1 - sample correlation. The reason for this was the amplitude of the spike often changed day to day, but the shape generally did not (the L^2^ norm is sensitive to changes in amplitude, but the sample correlation is not). Each paired distance was z-scored for normalization, and we calculated the dot product between the four distances and a weighting vector **w** to favor the most distinctive features, namely the spike time autocorrelation and interspike interval distribution,

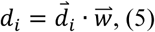

Where

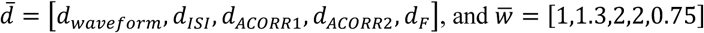

*d* was larger for neurons recorded on different channels than for neurons recorded on the same channel for both monkeys (**Supplemental Fig. 5c**). This difference likely reflects that some neurons recorded across days were the same. We performed complete-linkage hierarchical clustering on the set of *d* for each channel, and clusters were cut from the cluster tree using a threshold for a distance criteria of 4 (see **Supplemental Fig. 5b)**, which corresponded to *d* most likely to be observed between neurons on the same channel as compared with neurons on different channels (**Supplemental Fig. 5c**). Out of a total of 2671 neurons, the algorithm detected 1178 unique cells. **Supplemental Figure 6** shows how many neurons were recorded for a given number of sessions. **Supplemental Figure 7** shows channels with the most repeat recorded neurons identified by the algorithm.

### Permutation test for estimating statistical significance of repeated effects across sessions

For each condition (a-tDCS, c-tDCS, Sham), we calculated *ΔF* for all cells recorded more than once, and noted the percent of cells that were always modulated up or down past a given threshold *T*. We explored a range of *T* (from 0-10%) to avoid arbitrary parametrization. We tested the null hypothesis that neurons exhibited consistent firing rate changes by chance alone by relabeling the neurons within each condition to create 10,000 shuffled distributions. Thus, we calculated the probability that observed repeat *ΔF* occurred by chance given the measured *ΔF in that condition*, thereby accounting for differences in *ΔF* across conditions (for instance, the chance level of repeat increases during a-tDCS is higher than c-tDCS because more cells tended to increase their firing).

#### Torque Directionality

Polar plots of firing rate by torque direction were generated to visualize the pattern of spiking dependent upon the animal’s direction of wrist torque. To construct the rate map, torque direction was collected into bins of 6 degrees and the number of spikes in each bin was divided by the time the torque spent in that direction. To quantify the degree of directional selectivity, we calculated the mean resultant, *R*_*m*_, of the directional firing rate map:

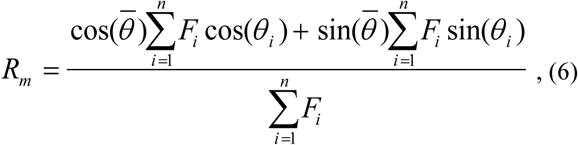

where 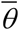 represents the preferred firing direction of the cell, and is calculated by

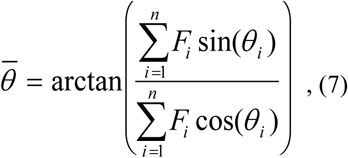

### Tuning rate functions

Firing rate maps of cell spiking in torque space were constructed by dividing the spike count within pixels of 2-dimensional torque data by the time spent by the cursor in that bin. Data were smoothed by a two-dimensional convolution with a pseudo-Gaussian kernel with a standard deviation of one pixel. Rate maps were compared across epochs using Pearson’s correlation coefficient, *ρ*, calculated by the MATLAB function *corrcoef*. The torque vector information for a given spike, *I* bits·spike^-1^, was inspired by Skaggs et al^47^, and is defined in equation 8:

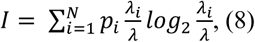

where the torque space was divided into N nonoverlapping pixels, *p*_*i*_ is the cursor occupancy probability of bin *i, λ*_*i*_ is the mean firing rate for bin *i*, and *λ* is the overall mean firing rate of the cell.

### Spike-triggered LFP and unitary spiking contribution to LFP

We calculated whitened spike triggered LFP (wst-LFP) features as described by Teleńczuk et al^58^. The wst-LFP technique applies spatial filters that decorrelate LFP signals in space, thereby eliminating features such as beta oscillations that are broadly distributed across the array. After this filtering (“whitening”) step, features in the spike-triggered averages are limited to electrodes adjacent to the triggering neuron, and peak at lag times that match predicted propagation speeds of axonal conduction. It appears that these features represent the unitary contribution of the single neuron to the LFP, reflected by post-synaptic currents following spikes.

We followed the same protocol as described previously^58^: we first calculated the covariance matrix (*C*) of the bandpass filtered, continuous LFP (15-300Hz, 3^rd^ order elliptic filter, results are similar with a Butterworth filter). Bad channels were removed by visual inspection from the analysis. Averages of the ongoing LFP about spikes (st-LFP; −50 to 50 ms) are calculated for each cell, and the wst-LFP were calculated by the matrix product of the st-LFP and a whitening matrix (*W*) derived from the covariance matrix:

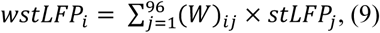

where

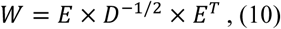

and *E* is the matrix of eigenvectors of the covariance matrix, and *D* is a diagonal matrix of eigenvalues.

To calculate the change in wst-LFP trough about spiking, we averaged all electrodes separated from the triggering neuron position by the same L_1_ (Manhattan) distance and took the minimum amplitude from lags 0-6ms relative to spike time. This window contains the full range of delays reported previously and observed in our data (**Figure 9b**).

### Neural population dynamics dimensionality reduction and analysis

Population spiking activity during the target tracking task was analyzed with a focus on neural dynamics during each cursor movement. **Supplemental Figure 8a** shows data for an example trial where time *t = 0* when the target appears on screen. The top panel of **Supplemental Figure 8a** shows the *FE*- and *RU-torques* registered by the manipulandum while the second panel shows a raster plot of spike times for all recorded neurons (*N = 49*) during that experiment. Similar to other studies^52^, the instantaneous firing rate of each neuron is approximated by convolving its spike train with a Gaussian filter with σ = 100 milliseconds and temporal resolution of *dt* =1 millisecond, as illustrated in the third panel in **Supplemental Figure 8a**. We verified that changing the definition of instantaneous rate did not change the qualitative nature of our results by testing one-sided Gaussian and Exponential filters, as well as spike counts in sliding windows of width σ ranging from 10-100ms. On each trial, the firing rate curve of every neuron was normalized to zero mean and unit variance (over the time course of the trial) to mitigate the biasing effects of heterogeneous spike counts across neurons and tDCS-induced changes. Neural population activity during trials was thus described by a time-dependent, *N*-dimensional rate vector, as depicted in the bottom panel of **Supplemental Figure 8a** (not normalized for illustration). The trial-averaged mean firing rate of a neuron was computed by averaging the rates at each time point across aligned trials. Target-conditioned mean rates were obtained by only including trials with the same target in the averaging. **Supplemental Figure 8b** shows the mean rate aligned on target 1, for two example neurons (one is up-modulated and the other is down-modulated). For all following analysis, only the activity taking place between *t* = 0 (target onset) and *t* = 0.5 seconds was considered. **Supplemental Figure 8c** shows the mean rates of all neurons (*N*_*cells*_ = 49) for each target.

Rate averaging was further conditioned on the four experimental epochs (Pre, tDCS/Sham, Post). We performed a principal component analysis (PCA) of neural dynamics for each epoch by concatenating all target-conditioned mean rates vectors into a large set of *N*_*cell*_-dimensional vectors. This procedure is a version of demixed PCA^54,55^ and extracts relevant subspaces where population coding takes place by taking task parameters into account. **Supplemental Figure 9** illustrates the results of this analysis for three experimental sessions with different tDCS modalities (sham, a-tDCS, c-tDCS) where we plot the projection of the target-conditioned population rates in the space spanned by the first two principal components (PC), alongside the target-conditioned mean torques for the corresponding tDCS epochs (**Supplemental Fig 9a**). Thus, for each epoch, a different PCA model was computed. Even if the shape of projected activity differed from one model to the next, the first two PCs were often sufficient to decode direction of motion for any tDCS modality since the task has two degrees of freedom. To check this, we trained a linear classifier to predict the target from the single-trial activity projected in the first two demixed PCs and found that the average of the 80%-20% split, cross-validated performance was between 80% and 95% (close to monkey performance) for all sessions. Trials where the monkey failed to reach the appropriate target were included in this spot check.

We estimated the dimensionality of ensemble trajectories using previously reported methods^56,57^. The dimension *D* of a set of points in *N* dimensions is computed as:

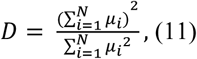

where *µ*_*i*_ represents the eigenvalues of the covariance matrix of the data used in the PCA model. If the *N* coordinates are statistically independent from one another, *D*=*N*, and if they are perfectly correlated, *D*=1.

Beyond the contribution of neurons to coding subspaces, we investigated the geometry of subspaces themselves. PCA produces linear subspaces (hyperplanes) whose orientation can be measured by principal angles. To measure how similarly oriented are two *d*-dimensional PC subspaces, we measured the *d* principal angles between them {*θ*_1,_…, *θ*_*d*_} and define their ***orientation similarity***, *S*_*ori*_, as the mean cosine of these angles: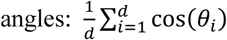. Identical subspace will have *S*_*ori*_ = 1, while orthogonal spaces will have *S*_*ori*_ = 0. In a similar analysis, we quantified how individual neurons contributed to a subspace spanned by a given set of PCs using a measure we call the participation score ***(Supplemental Methods)***. This analysis, shown in **Supplemental Figure 9b**, was consistent with findings reported in Figure 8e.

We computed the three comparative quantities described above, (i) dimensionality difference, *ΔD*, (ii) correlation of participation score, *C*_*part*_, and (iii) orientation similarity, *S*_*ori*_, for three tDCS epoch pair combinations (*Pre-Stim, Pre-Post)* across experiments with different tDCS current doses (low dose ≤1mA and high dose >1mA).. Comparison between tDCS and Sham experiments were made using independent t-tests. The number of neurons *N* varied from session to session and we discarded any session with *N<*10 (experiments included; Monkey S: 59, Monkey W: 40), with *N* ranging from 10 to 54 with median of 27. Results for both monkeys were similar, and we combined experiments to increase power for statistical analysis. **Supplemental Table 1** further breaks down the number of samples by session type. **Supplemental Figure 10** shows box plots of all sampled quantities along with these p-values.

## Data availability statement

The data that support the findings of this study are available from the corresponding author upon reasonable request.

## Code availability statement

The code that supports the findings of this study are available from the corresponding author upon reasonable request.

## End Notes

## Acknowledgements

We thank Richy Yun, Mihály Vöröslakos, Steve I. Perlmutter, Marco Capogrosso, and Brian J. Mogen for their invaluable consultations during the preparation of the manuscript, Rebekah Schaefer for animal care, and Larry Shupe for technical assistance. This work was supported by EEC-1028725 (ARB), F30NS100253 (ARB), R01NS12542 (EEF) and the NSF ERC Center for Neurotechnology

## Author contributions

A.R.B, S.Z., and E. E. F. conceived the project. A.R.B. and S.Z. performed surgeries. A.R.B., H.B., and A.M. performed the experiments. G.L. performed the analyses underlying Figure 8 and Supplementary Figures 8-10, and A.R.B performed all other analyses. A.R.B, E.E.F, and G.L. wrote the paper.

