## Supplemental Materials for "Cortical network mechanisms of anodal and cathodal transcranial direct current stimulation in awake primates"

**The PDF file includes:**

Supplementary Figures

Supplementary Tables

Supplementary Discussion

Supplementary References

**Supplemental figure 1.** Stable delivery of tDCS by stimulator. **(a)** Testing circuit. We measured the voltage drop across a 1 kOhm resister because the input range of the measurement device was ±5 V. The simulated tissue/electrode impedance was 14kOhms, which was greater than experimental conditions. **(b)** Constant current was stabley delivered for 25 minutes. At this load, the stimulator was delivering current corresponding to its compliance voltage.

**Supplemental figure 2.** Task performance and behavior during tDCS and Sham. Each plot shows histogram of changes for a given performance metric from Pre to Stim/Sham, with the median indicated along the x-axis with an 'X' (Sham: light gray, a-tDCS: red, c-tDCS:black). There was no difference between the time it took monkeys to complete trials (trial duration, $p>0.05$, **a**), or the time it took monkeys to move the cursor to the target following the variable hold period (response time, $p>0.05$, **b**). There was a small but significant difference between the changes observed during tDCS and Sham for the median torque produced during trials (**c,** median Δtorque, Sham=-2.9e-3 in·lbs, a-tDCS = 5.7e-3, c-tDCS = 5.2e-3) and number of successful trials per minute (**d**, trials with duration <6secs, Sham=-0.08 trials/min, a-tDCS =-0.72, c-tDCS =-0.6). Overall, behavior was tightly controlled during tDCS and was similar across experiments, because the monkeys were overtrained and completed the task at peak performance. Regular performance was important to eliminate any potential confounds associated with behavioral changes.

**Supplemental figure 3.** Average change in spiking for every experiment. Data plotted is mean ± standard error, and the number of neurons recorded is indicated in parentheses. Blue data points: c-tDCS, red data points: a-tDCS.

**Supplemental figure 4.** Population composition of FS and RS cells. Both monkeys had comparable ratios of RS:FS cells. Top: distribution of waveform width across all cells recorded in each monkey. Average waveform and standard error for FS (green) and RS (orange) cells shown at bottom.

**Supplemental figure 5.** Algorithm for longitudinal cell identification. **(a)** Five statistics of firing were calculated for each neuron during Pre, including average waveform shape, inter-spike interval histogram, spike-time autocorrelogram at two timescales, and average firing rate. We calculated pairwise differences for each metric across all recorded neurons, z-scored difference measure, and weighted the combined difference vector **d** by **w** to weight certain factors, such as autocorrelation, more heavily. **(b)** Example clustering results for a single recording channel over many experiments. We performed complete-linkage hierarchical clustering on the weighted set of *d* for each channel, using a distance threshold criteria of 4 derived from distributions shown in **(c)**. **(c)** Distribution of *d* between neurons recorded on the same channel (blue line) and neurons recorded on different channels (red line) in Monkey S **(i)** and Monkey W **(ii)**. There is a higher frequency of low *d* between neurons recorded on the same channel, reflecting the fact that some neurons are repeatedly recorded during separate experiments. We derived the cluster threshold cutoff using these two distributions, selecting a value (4) that corresponded to the *d* most likely to be observed between neurons on the same channel as compared with neurons on different channels.

**Supplemental figure 6.** The number of cells recorded across sessions was similar for all conditions as detected by longitudinal cell tracking algorithm. Most neurons were recorded only once for a given condition (N=1162), but each condition (a-tDCS, c-tDCS, Sham) had a sizable number of neurons recorded more than once. These neurons permitted estimation of the reliability of tDCS effects in single neurons across sessions.

**Supplemental figure 7.** Results of longitudinal cell identification algorithm. Exemplary channels with unusually high repeat neurons are shown. Each neuron is represented by four statistics (Scale bars in top left. From top to bottom: average waveform, average firing rate (grey circle), inter-spike interval histogram, spike time autocorrelation.) Neurons identified across sessions are plotted together in boxes.

**Supplemental figure 8.** Task-specific population dynamics. **(a)** Example dynamic quantities for a target tracking task (N=49 neurons). Target onset occurs at t=0 (red bar on time axis). Top panel:

manipulandum x- and y-torque trajectories for this target. Second panel: population raster plot, each dot represents a spike time. Third panel: spike times of single neuron (black dots) and Gaussian-filtered rate ($\sigma$=100 milliseconds). Bottom panel: population firing rate curves as in third panel, colors match those of second panel (not normalized for illustration). **(b)** Target-specific, cross-trial spiking for two example neurons. Top: cross-trial raster plot. Bottom: mean firing rate curve. Shaded area shows one standard error of the mean. **(c)** Population mean firing rate curves (each color indicates a specific neuron) separated by target number for same time interval as (b).

**Supplemental figure 9.** Examples of ensemble trajectories in 2D manifold. PCA–related quantities for three representative sessions. **(Left)** Sham, N=29. **(Middle)** a-tDCS (2 mA), N=45. **(Right)** c-tDCS (3 mA), N=49. For all panels: Target-specific, averaged population firing rates projected in first two PCs (left columns) and averaged manipulandum torques (right columns). Each row indicates tDCS epoch (Pre, Stim, Post). Colors indicate target identity, circle markers indicate t=0 seconds (target onset), square markers indicate t=0.5 seconds. **(b)** *Participation scores* of all neurons, for subspaces spanned by first two PCs (d=2) of PCA models PRE, STIM and POST, respectively. Neurons are ordered in decreasing order of score in model PRE. Dotted line indicates score that equally participating neurons would hold.

**Supplemental figure 10.** Statistics of population coding metrics. **(a)** Change in dimensionality of ensemble trajectories. Dimensionality is normalized by Pre to document differences across experiments. Main Figure 6 shows pre-normalization values. **(b)** Correlation coefficient for participation score of neurons to the subspace spanned by the first two (d=2) PCs of PCA models. **(c)** Orientation similarity measure between subspace spanned by first two PCs (d=2) of PCA models. For all panels: Box edges show first and third quartiles, internal bar shows mean, whiskers show extremal values. Comparative quantities are plotted for model pairs. From left to right: Stim-Pre, Post-Pre. In each plot, quantities are shown for tDCS low ($\leq$ 1mA, left) and high (>1mA, right) doses, for three conditions: sham (white), a-tDCS (red), c-tDCS (black). Sham box is repeated for low and high doses. P-values are computed from comparison with sham samples with an independent t-test. Effects during Post reflect activity for 30 minutes after tDCS was turned off.

| | a-tDCS | c-tDCS | Sham |
|---|---|---|---|
| low dose | n=14 | n=14 | n=40 |
| high dose | n=13 | n=18 | |

**Supplemental Table 1. Number of experiments for each population dynamics analysis**

| *Metric during PRE epoch* | *p-value*** *(group medians)* |
|---|---|
| Firing rate (Figs 1 and 2) * | 0.99 (1.97 1.88 2.40 2.21 2.83) |
| Firing rate during "contracting" phase (Fig. 3) † | 0.90 (2.16 1.93 2.30) |
| Firing rate during "resting" phase (Fig. 3) † | 0.97 (1.59 1.82 2.05) |
| Directional tuning strength ($R_L$) * | 0.16 (0.08, 0.08, 0.07, 0.08, 0.07) |
| Spike Information (Fig. 6) * | 0.36 (0.24, 0.27, 0.23, 0.24, 0.25) |
| Dimensionality (D) of population activity (Fig. 7) | 0.49 (3.77, 3.22, 4.20, 3.58, 3.82) |
| ws-LFP amplitude (Fig. 8) † | 0.66 (-0.08, -0.08, -0.08) |

* for groups Sham, LD a-tDCS, HD a-tDCS, LD c-tDCS, HD c-tDCS

† for groups Sham, a-tDCS, c-tDCS

** one-way ANOVA

**Supplemental Table 2. Equivalence of baseline values before intervention**

**Supplementary Discussion**

While we tried to match human tDCS as closely as possible, some differences were unavoidable: to record intracortically requires a bone defect where the recording electrode wire bundle passes into the intracranial space, and we used a thin titanium strap to hold the bone flap in place immediately post implantation. Implants and boney defects may distort the electric field or shunt current. On the other hand, we found that the bone flap reintegrated fully, the residual passage for the wire bundle was very small, and extracranial tissue recovered fully. Furthermore, the skull itself is naturally porous due to features like cranial sutures and Haversian canals, and the relatively snug passage for the wire bundle may not be so different.

Some potentially important differences between humans and monkeys include the size and distance of the electrodes, shape of the head (in particular the brow), potential differences in cortical neuron subtypes, and the amount of current that passes through the brain, which depends on the tissue layers and the effects of implanted material, and hair is often left in place during human experiments, whereas we removed it. We could not experimentally measure the intracranial electric field induced by scalp stimulation as done in recent preparations with humans and monkeys[1–4], because this requires that electrodes are placed along the gradient expected between the stimulating electrodes. Electrodes of the Utah array, on the other hand, are orientated tangentially to the cortical surface, which maximizes the number of neurons recorded, but are limited for estimating the currents induced in the brain by tDCS. Moreover, the electrode tip capacitance does not allow accurate measures of DC potentials.

Thus, due to technical constraints, we have not reported estimations of the electric field (see also[5]). Luckily, multiple reports exist across multiple montages and currents in humans and monkeys, and find that current density at the skin is a reasonable surrogate for intracranial electric field (see main text, discussion). From the literature we note that electric fields evoked in the brain by scalp stimulation of a given current density are within an order of magnitude across different subjects, and between monkeys and humans. We spanned a 16x range of current density in our study, which by inference from past reports, should result in field strengths ranging from roughly 0.015 - 1 $V \cdot m^{-1}$. Considering that human tDCS produces fields of about 0.2-0.8 $V \cdot m^{-1}$, it is likely that our stimulation intensities produced field intensities that overlap with those induced by human tDCS. It is worth noting that a recent human trial tested a range of currents outside of normal tDCS, and the lowest current density that produced changes (in EEG alpha oscillations) was similar to that which produced significant single cell effects in our study[2] ($0.125 mA/cm^2$ versus $0.11 mA/cm^2$). We confirmed that tDCS intensity is a critical parameter – larger currents produced larger effects in disparate measures such as single cell firing rates, rate-normalized population dynamics, and the wst-LFP. We found effects in the primate brain at current densities within clinical safety limits, but at currents that were about four times higher than those generally applied to humans ($0.11 mA \cdot cm^{-2}$). Should currents this great be required in humans, comparable intensities have been applied to the scalp of awake humans subjects, albeit with some side effects such as headaches, nausea, sleepiness, and tremor. In particular, between 1960 and 1998, a few studies[6] delivered tDCS through much smaller scalp electrodes with current densities of $0.2 mA \cdot cm^{-2}$, well within the range of effects in our study. One older study successfully applied 2.4 $mA/cm^2$ to one patient with local anesthetic under the electrode[7].

**Supplementary Methods**

To quantify how individual neurons contributed to a subspace spanned by a given set of PCs, we defined the neuron's ***participation score***: the norm of the canonical neural vector $v_i$ (*N*-dimensional vector with zeros everywhere but at the $i^{th}$ position) linearly projected in the subspace. **Supplemental Figure 9b** shows the participation score of all *N* neurons for the space spanned by the first two PCs in each epoch-specific PCA models (neurons are ordered by their participation score in the *PRE* model). Some neurons were more informative than others, and the spread in participation scores reflects that. In contrast, if all neurons contributed equally to the subspace, their scores would all be $\sqrt{d/N}$ where $d$ is the dimension of the subspace ($1 \leq d \leq N$, see dashed line in **Supplemental Figure 9b**). For a given subspace dimension $d$, we measured the difference in participation scores between two epoch-specific PCA models by computing the Pearson correlation coefficient, $C_{part}$, of the pairs $\{(s(d)_i^a, s(d)_i^b)\}_1^N$ where $s(d)_i^a$ denotes the participation score of neuron *i* in the first *d*-dimensions of PCA model *a*. A high correlation coefficient indicates that neurons have similar participation scores in models *a* and *b*, while a low coefficient indicates a big change in participation. Variability in participation score is expected between epochs, even for sham sessions where no stimulation is administered since neural activity is noisy and drifts during the course of the experiment. This is especially true for $d \ll N$ since low-dimensional subspaces have many "free degrees of freedom" to move in (i.e. *N*-*d*). Nevertheless, we concentrate on *d*=2 since the task itself involves two degrees of freedom (i.e *x*- and *y*-torques), a fact that should be recovered in task-relevant neural activity. We find that tDCS stimulation can induce greater-than-normal variability in participation score for *d*=2, indicating a rearrangement of neurons' roles in supporting task-relevant subspace (see statistical tests reported in **Supplemental Figure 10**). We verified that these results generally hold for *d* up to 5.
