## Supplementary figures and images for "Cortical network mechanisms of anodal and cathodal transcranial direct current stimulation in awake primates"

### Supplemental Figures

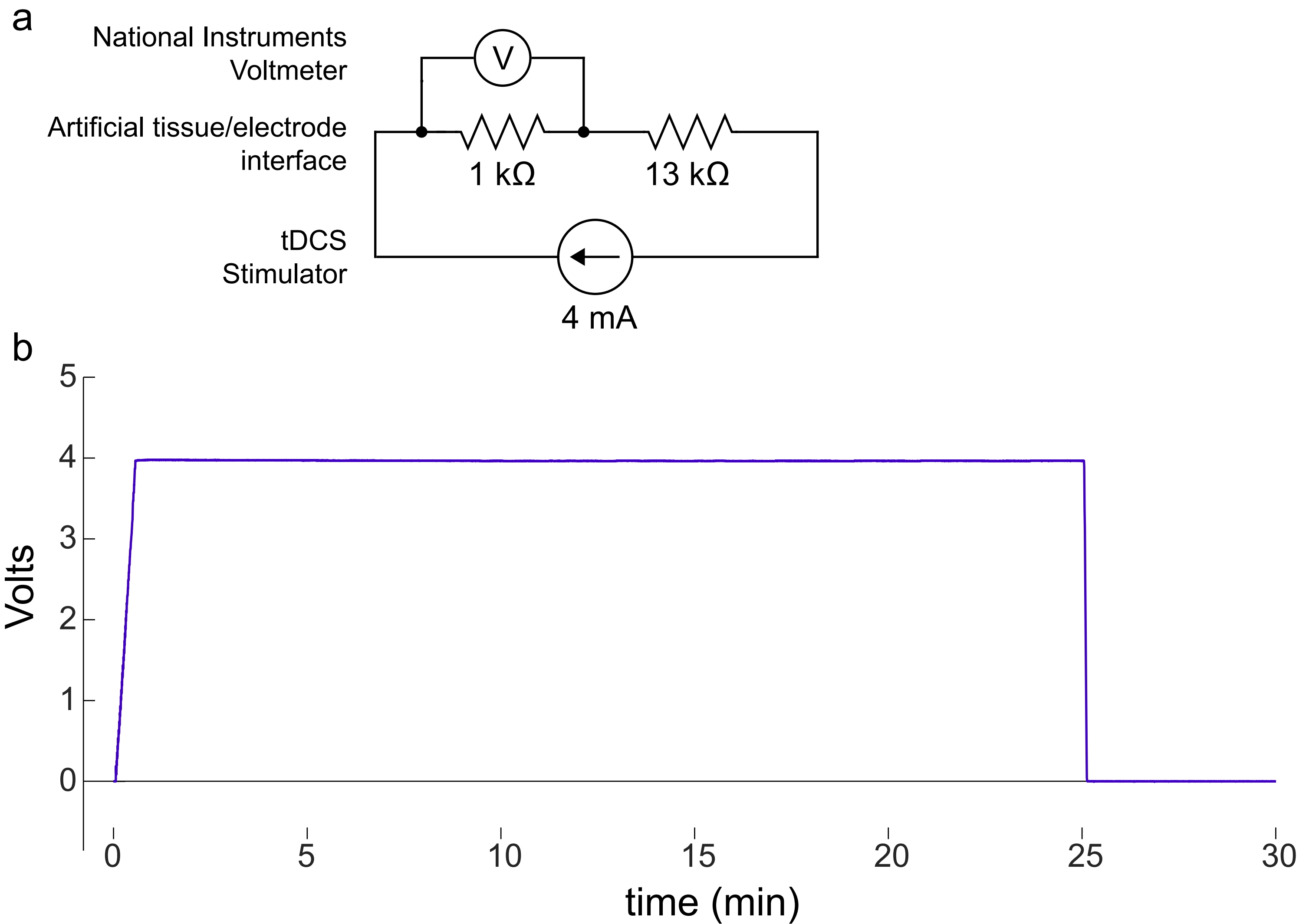

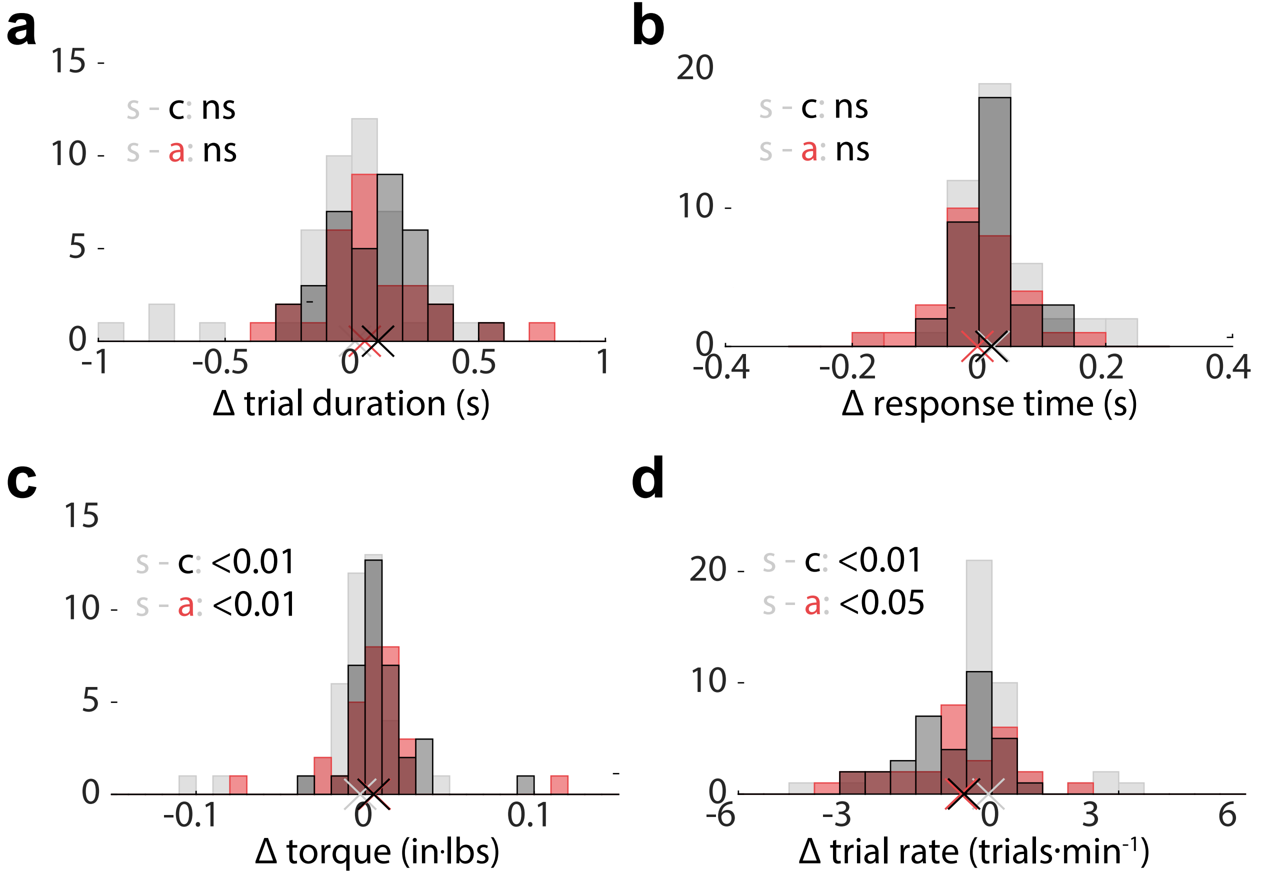

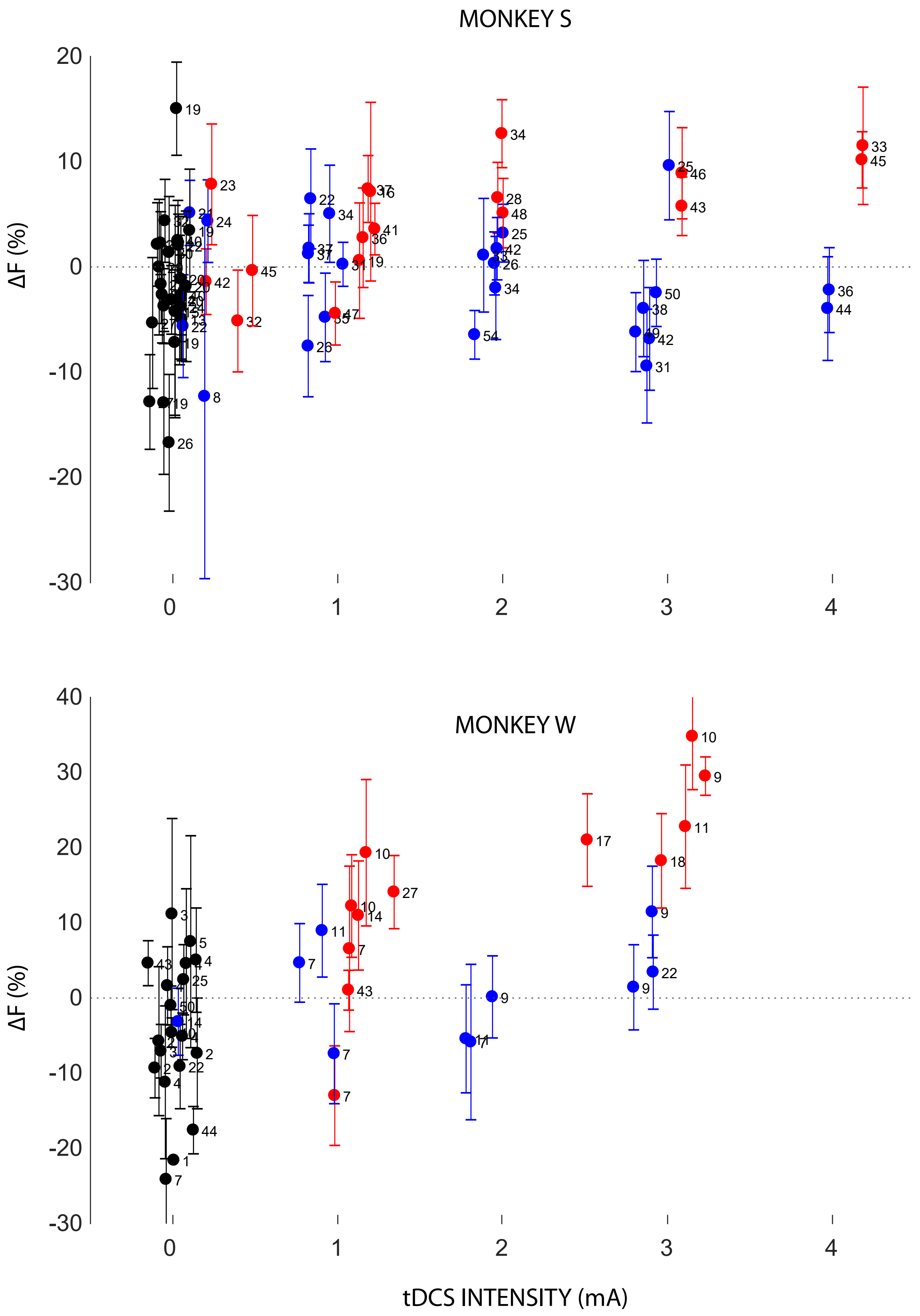

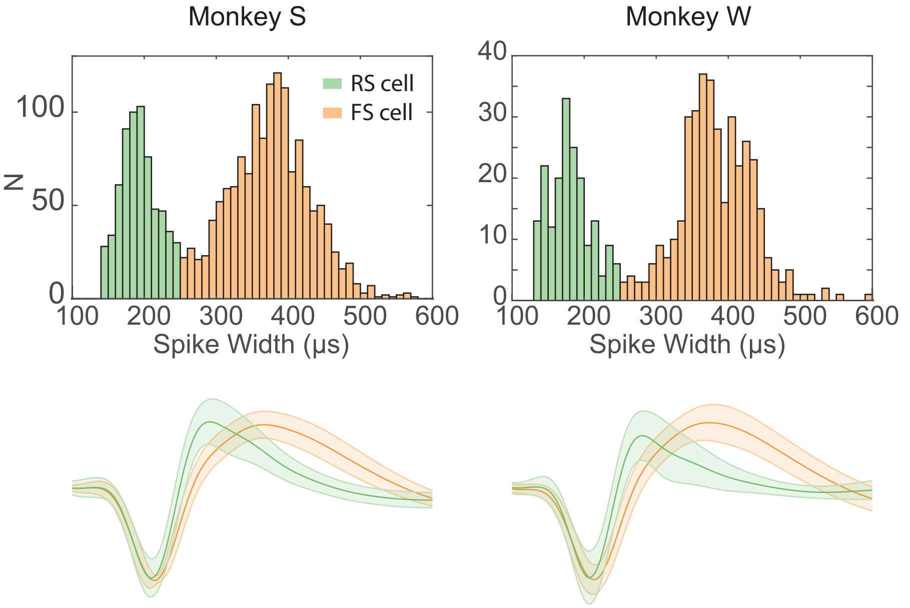

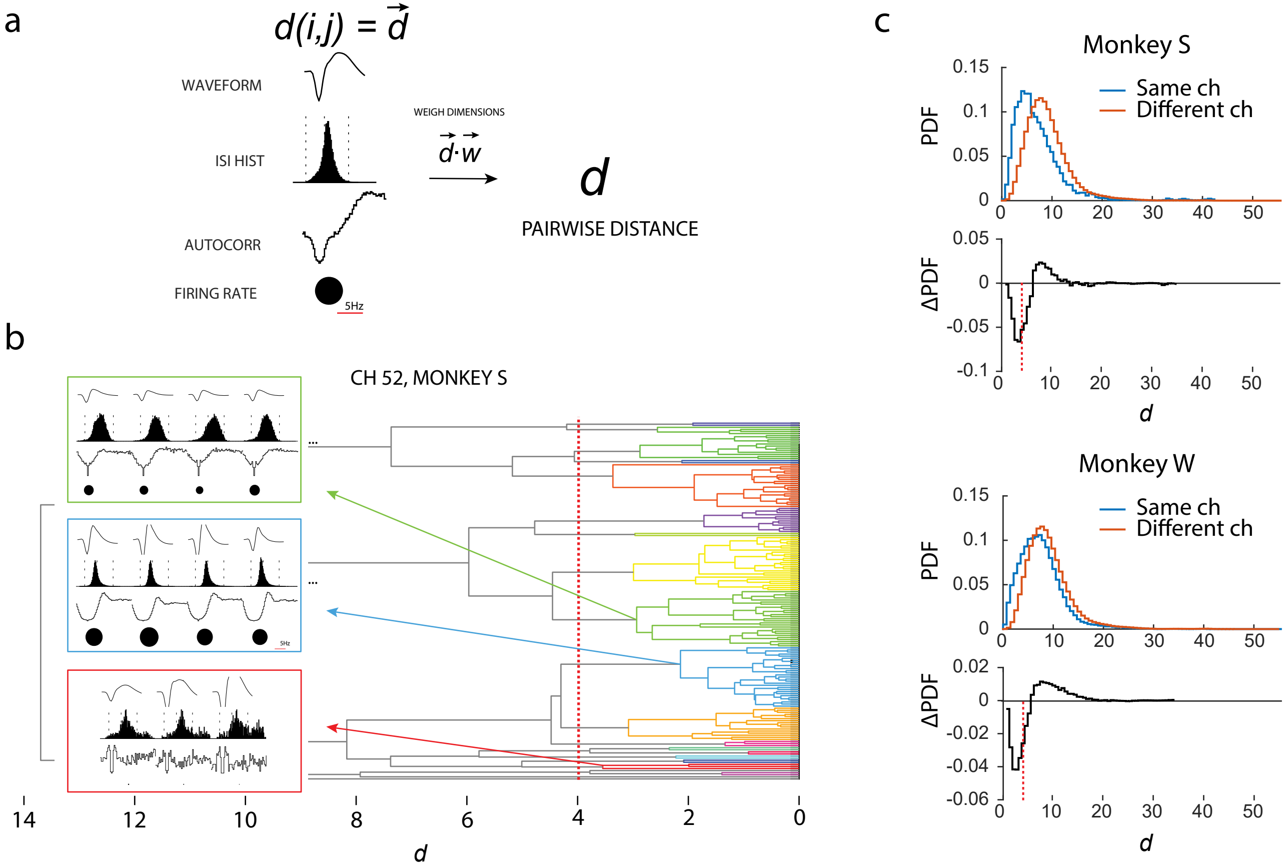

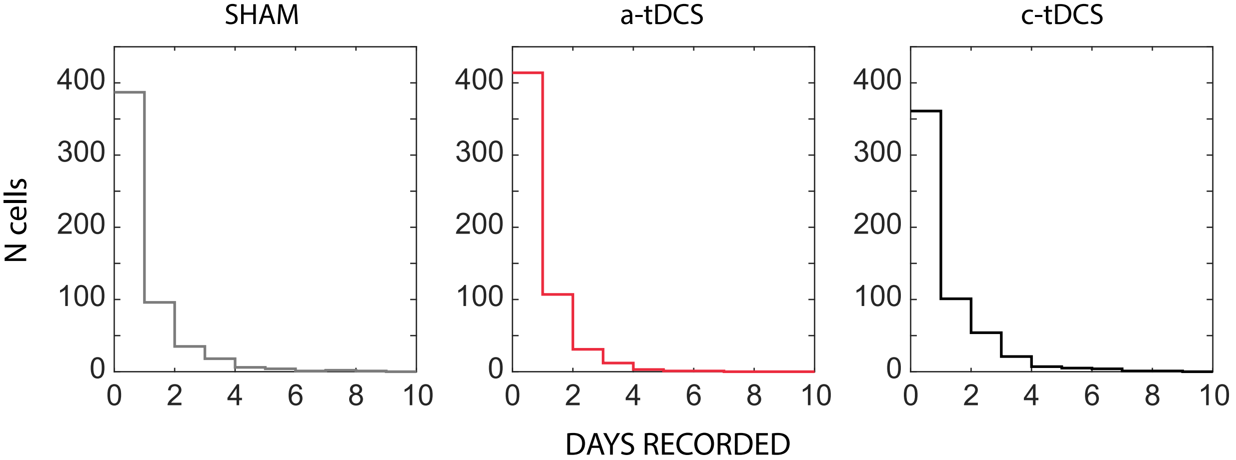

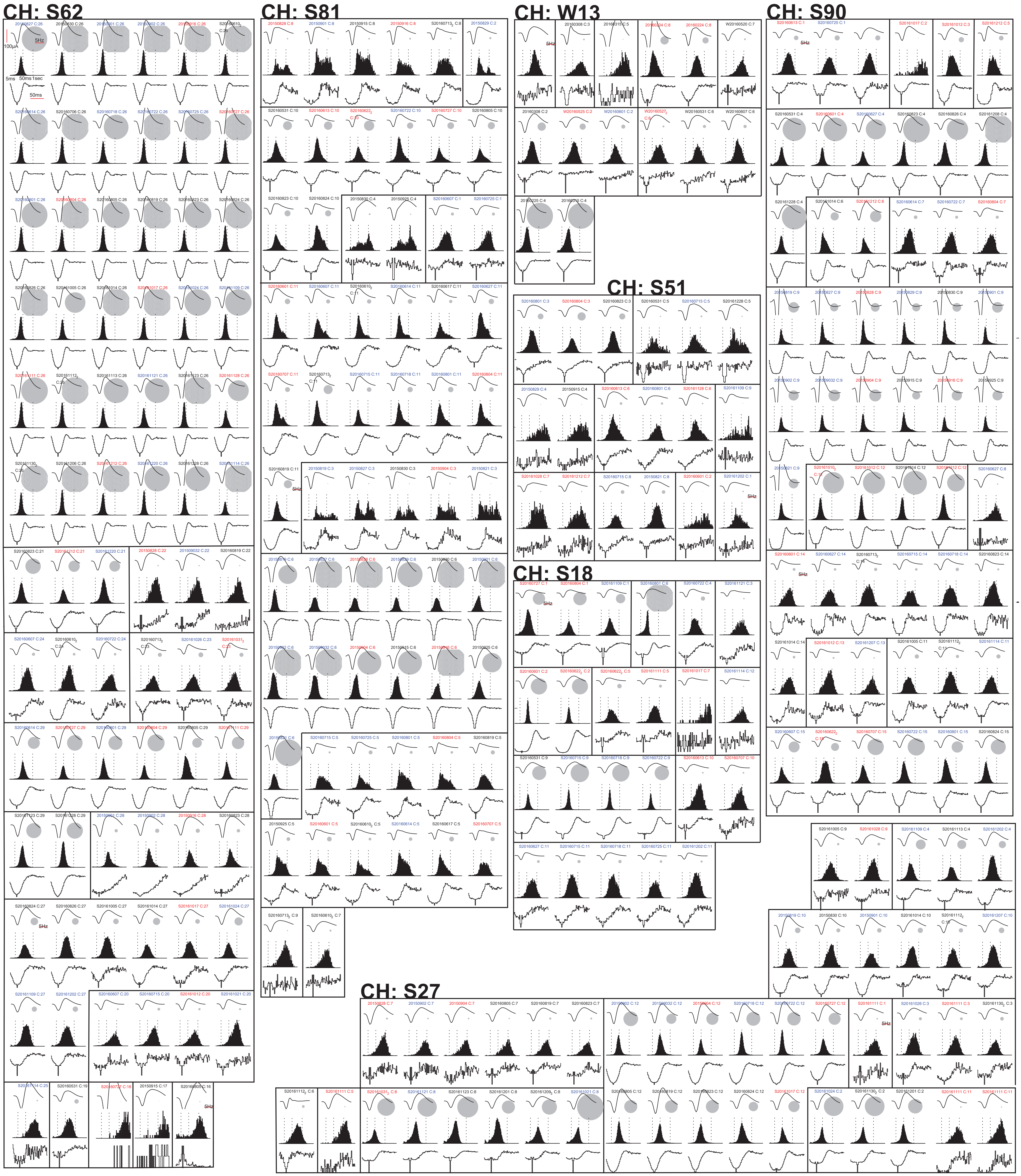

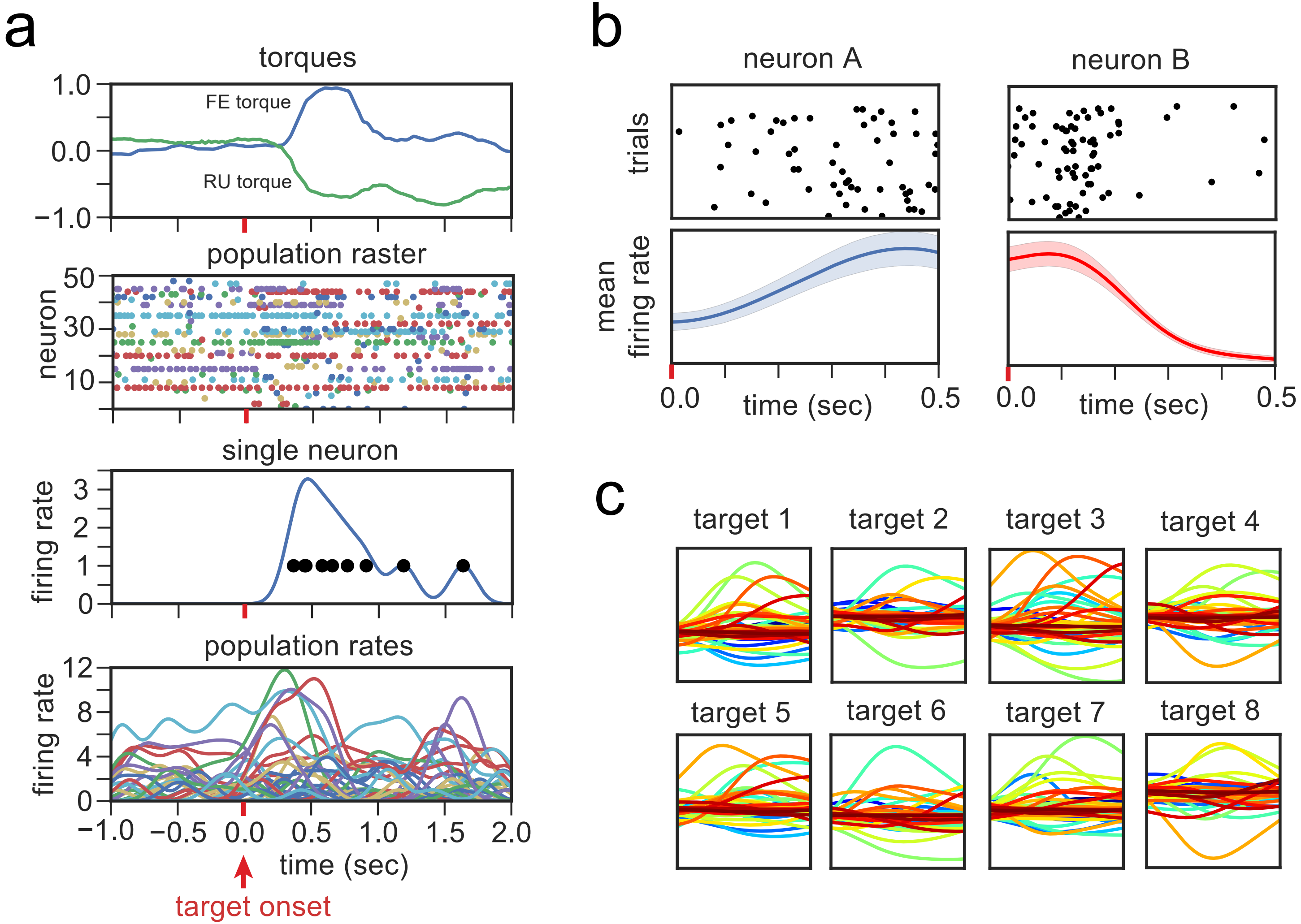

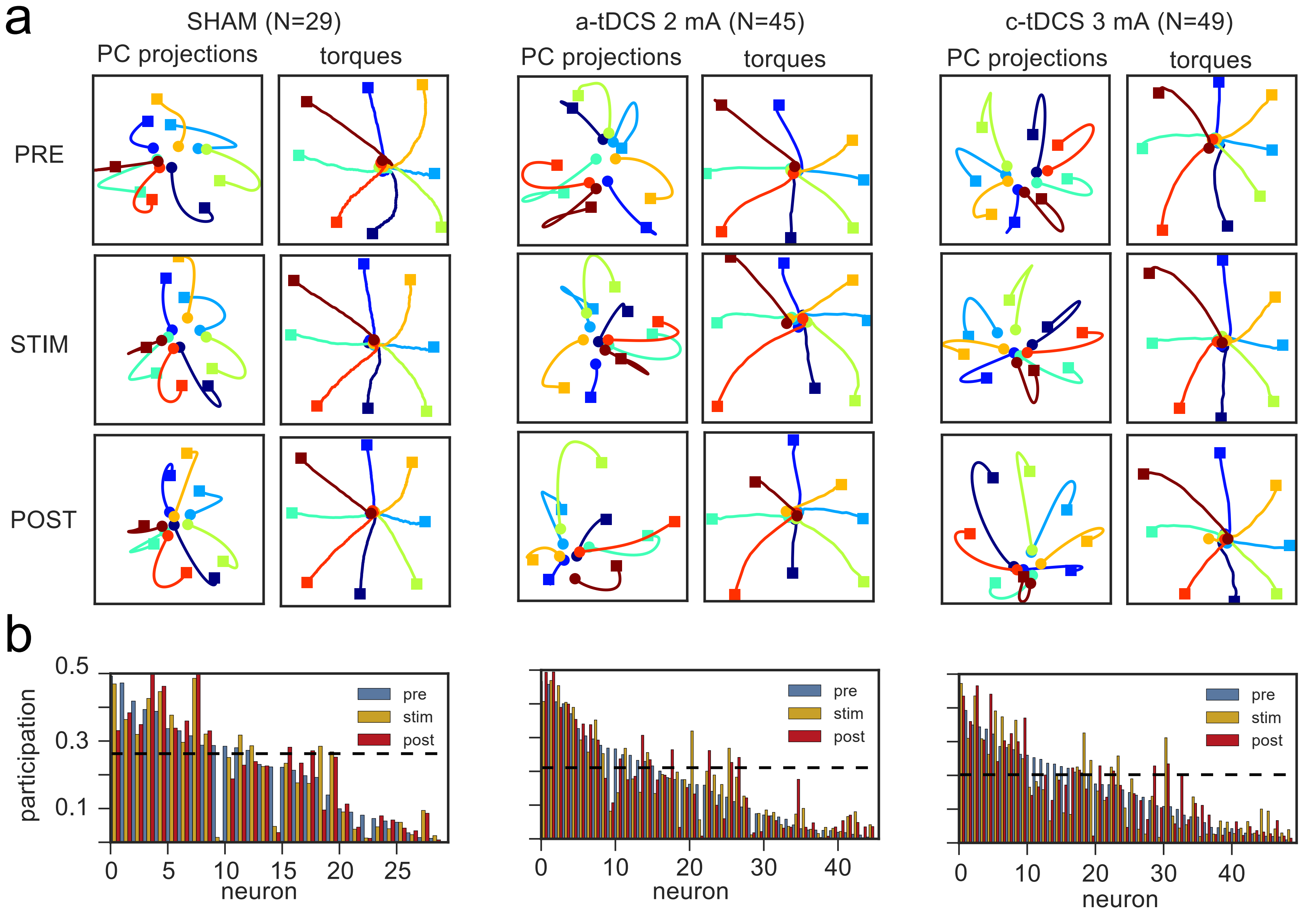

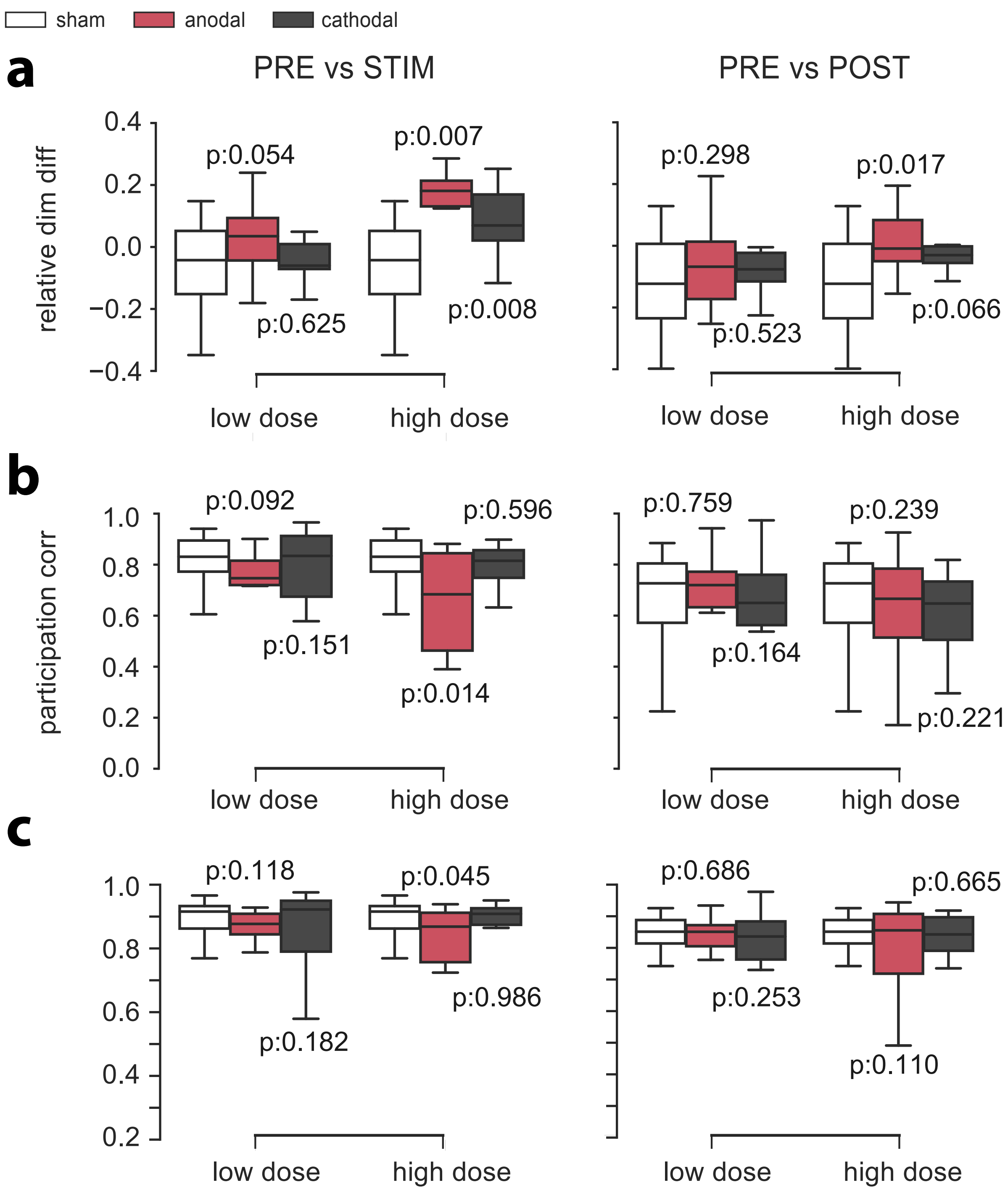
